# A consensus atlas of human brain development defines cell type-specific maturation trajectories across the lifespan

**DOI:** 10.64898/2026.07.16.738788

**Authors:** Sridevi Venkatesan, Patricia Nano, Jonathan Werner, Sonia Malaiya, Brian Herb, Yuan Gao, Li Wang, Aparna Bhaduri, Tomasz J. Nowakowski, Carlo Colantuoni, Seth A. Ament, Jesse Gillis

## Abstract

Human neurodevelopment is a continuous process that begins prenatally and extends into postnatal life. Current transcriptomic datasets are limited, fragmented across analytical frameworks, precluding comprehensive reconstruction of cellular trajectories linking developmental states to mature cell types. Here we present a consolidated cellular-resolution transcriptomic atlas of human brain development from the onset of neurogenesis to adulthood, covering ∼2.2 million cells from 156 donors across nine studies. All data were reprocessed from raw sequencing reads and annotated within a unified cell-type taxonomy, enabling reliable mapping across the lifespan. Cell types were highly replicable across heterogeneous datasets, enabling us to chart their maturation and cortical layer localization over time. We identify dynamic gene programs predictive of cell-type maturation, validate gene modules tracking known fate transitions, and leverage our atlas’ scale to characterize rare populations, including microglia. This resource establishes a standardized reference of human brain development and maturation gene modules for future comparisons across model systems, species, and disease states.

## Introduction

Human brain development is a protracted and continuous process spanning pre- and post-natal life, encompassing progenitor expansion and differentiation, the generation of a wide spectrum of neuronal and glial cell types, and the progressive maturation of these populations into stable adult identities. Single-cell transcriptomic atlasing efforts have substantially advanced our understanding of this process, revealing the molecular programs that drive developmental transitions from progenitors to maturing cell types^1–10^. However, existing datasets are drawn from distinct developmental stages, use different technologies (single-cell vs nucleus), and are processed heterogeneously, resulting in a fragmented view of a fundamentally continuous process.

Early bulk transcriptomic studies of human brains identified a major shift in gene expression at the transition from prenatal to postnatal life^11,12^. Single-cell studies have further shown that features of mature neuronal function emerge after birth in most cell types^6^ and characterize cell type-specific maturation dynamics in detail. But the timepoints and analytical pipelines used to study these dynamics vary across studies, leading to differing conclusions on how cellular identity is refined over time and the similarities across prenatal and adult timepoints^13–15^. Unlike mouse studies where dense sampling of brains across timepoints is more tractable^16–18^, human data is inherently fragmented. As a result, it remains unclear when developing neurons acquire stable adult transcriptional states, what molecular changes underlie cell type-specific maturation, and how these dynamics vary across cell types.

Resolving this challenge requires rigorous re-assessment of independent datasets to annotate replicable cell types^19^ that enable coherent evaluation of maturation trajectories. Integrative analyses provide consistent cell type nomenclature and serve as a reference for comparative studies, seen in previous work from the Human Cell Atlas^20,21^ and BRAIN Initiative Cell Census Network^22^. Efforts to improve *in vitro* neural organoid models, cross-species developmental comparisons, and understanding early origins of disease all require a reliable reference of *in vivo* neurodevelopment. This necessitates integrating single-cell data across the full span of human development with an analytical framework robust to study-specific artifacts.

We present a unified single-cell and single-nucleus transcriptomic atlas of human brain development from gestational week 6 to adulthood, covering 2.18 million cells from 156 donors across nine studies. All data were reprocessed from raw sequencing reads and cell types were annotated using pre-defined meta-analytic marker sets ^23,24^. We verify the replicability of 35 cell types^25^ across time points, studies, and technologies, delineating critical periods when rapid maturation towards adult identity is completed. Projection of our atlas onto spatial transcriptomics datasets^26^ recapitulates the inside-out sequence of cortical layer formation, indicating that gene expression programs characteristic of adult cell types are detectable early in developing neurons.

We further identify gene sets predictive of cell type maturation^27^, revealing established signatures such as the neurogenesis to gliogenesis switch in progenitors and increase in marker expression across cell types. Our atlas enables the identification of continuous maturation trajectories even in rare populations such as microglia. Finally, we show that expression patterns of gene modules specific to early neuron and cortical layer identity^28^ are recapitulated in our atlas, and we extract temporally dynamic gene modules^29^ that are co-expressed both within and across cell-types revealing multi-level gene regulation^30^ in the developing brain. These results show the advantage of our uniformly processed atlas in defining multiple axes of neural maturation.

Our work converts fragmented temporal sampling of single-cell datasets into a continuous view of neurodevelopment, defining cell type-specific maturation trajectories across the lifespan. Our results are available to explore in an interactive webserver (<u>brain-development browser</u>) and the underlying data will be available via CellxGene. This comprehensive resource will serve as a reference for future experimental and comparative studies of brain development.

## Results

### Uniform processing and annotation strategy to define cell types across lifespan

We processed raw FASTQ files from 9 recently published single-cell and single-nucleus datasets^1–9^ to obtain an integrative atlas of the developing human brain. After quality control for minimum number of genes, mitochondrial reads, and doublet filtering, we obtained 2,157,582 cells across 156 donors, spanning first trimester to adulthood (**Fig 1a**). Cortical regions, forebrain, and ganglionic eminences dominated our atlas as these regions have been the most extensively sampled in previous studies.

**Figure 1:**
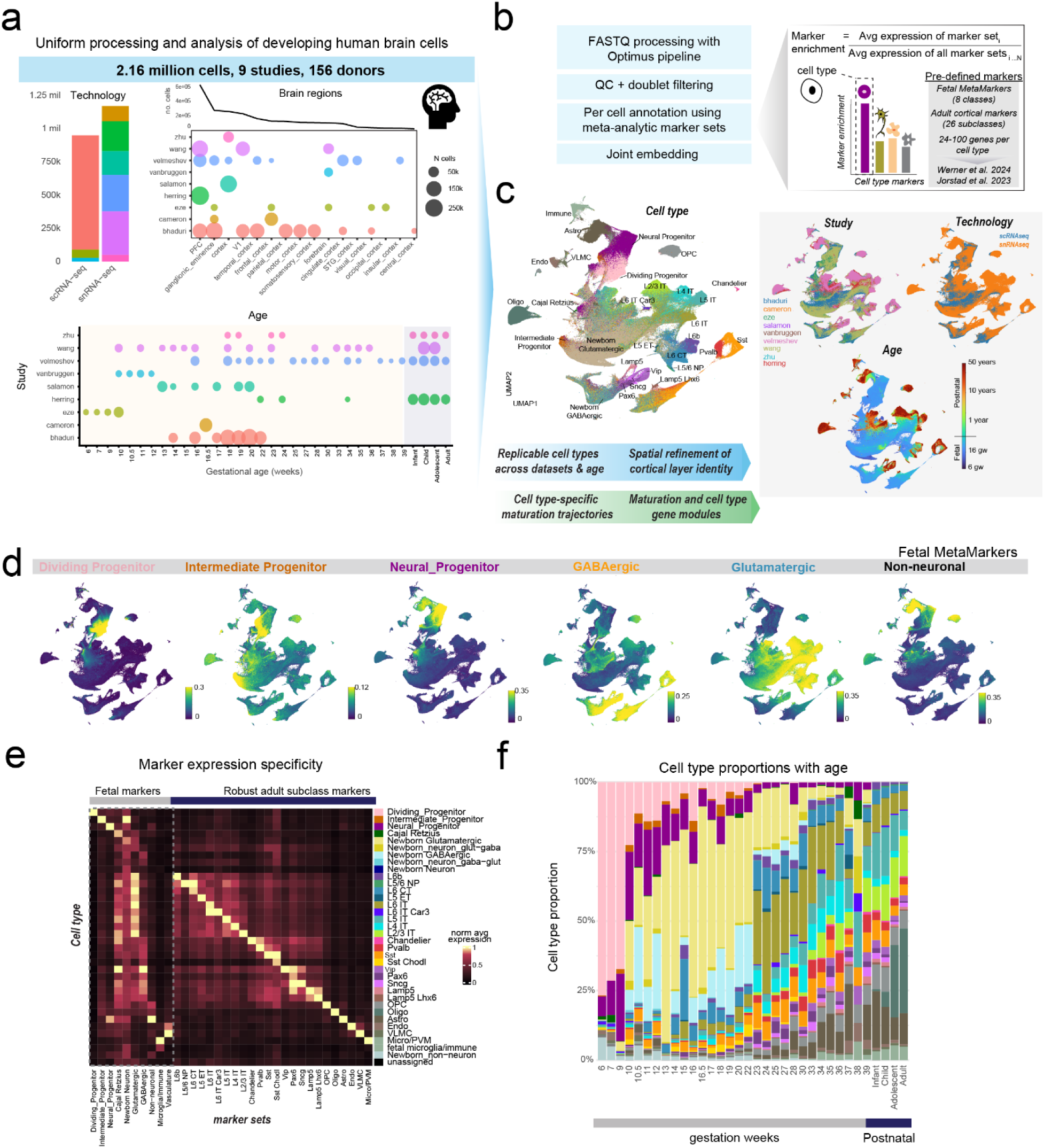
Pipeline for uniformly processed atlas of human brain development across lifespan. **(a)** Technology, time points, and brain regions sampled from 9 studies in our uniformly processed atlas. Single-cell and single-nucleus transcriptomic data from cortical regions, forebrain, and ganglionic eminences across human lifespan (1st trimester to adulthood) were uniformly processed. **(b)** FASTQ files were processed using the Optimus WARP pipeline and cells passing QC filters were annotated at the single-cell level using meta-analytic cell type marker sets (24-100 genes each) by computing a per-cell marker enrichment score. Each cell is assigned the cell type with highest marker enrichment. **(c)** UMAP plots show consistent grouping of cells by cell type across 34 subclasses, spanning timepoints, technologies and datasets. UMAPS were generated using CONCORD embeddings. **(d)** UMAP plots show aggregate expression of meta-analytic marker sets for fetal brain cell types. **(e)** Heatmap shows average expression of marker sets in each of the annotated cell types across the atlas. **(f)** Proportion of cell types at each time point.

To address the challenge of uniformly annotating cell types across diverse developmental timepoints, datasets, and technologies, we relied on predefined marker sets for 24 subclasses from an adult cross-regional consensus taxonomy ^23^ and progenitor and newborn neuron markers from meta-analyses of the prenatal brain^24,29^. We annotated cell types at the single-cell level based on highest enrichment for the meta-analytic marker sets^31^ (**Supplementary table S1**) and reached a consensus between the prenatal and adult annotations to obtain 35 cell types in total (**Fig 1d-e**). These include three progenitor subclasses: Neural progenitors (expressing canonical radial glia markers *VIM*, *HOPX*^32,33^, **Supplementary Fig 1**), Intermediate Progenitors (expressing *EOMES*), and Dividing progenitors (expressing *MKI67*); Cajal Retzius cells, Newborn Glutamatergic, Newborn GABAergic neurons; a small population of newborn neurons with mixed glutamatergic and GABAergic identity (newborn_neuron glut-gaba, newborn_neuron gaba-glut biased towards glutamatergic and GABAergic identity respectively); mature Glutamatergic subclasses: L6b, L5/6 NP, L6 CT, L5 ET, L6 IT Car3, L6 IT, L5 IT, L4 IT, L2/3 IT, CT-corticothalamic, IT-intratelencephalic, ET-extratelencephalic; mature GABAergic subclasses: Pvalb, Chandelier, Sst, Sst Chodl, Vip, Sncg, Pax6, Lamp5, Lamp5 Lhx6; Glial and non-neuronal cell types: Astrocytes, Oligodendrocyte Progenitor Cells (OPCs), Oligodendrocytes, Vascular Leptomeningial cells (VLMC), Endothelial, Immune cells, and newborn non-neurons.

Even with our unsupervised per-cell annotation approach that is distinct from typical de novo clustering-based annotation, cells of the same type clustered together in the UMAP across datasets and technologies (**Fig 1c**). Aggregate expression of marker sets accurately identified the corresponding cell types (**Fig 1e**, average AUROC for marker expression specificity = 0.96 across 28 celltypes, excluding transitionary newborn celltypes). Cell type proportions changed over age following expected patterns, with initial expansion of the progenitor pool in GW6-9, then the emergence of newborn glutamatergic and GABAergic neurons from GW10-20, followed by an increase in deep layer-specific glutamatergic neurons and finally upper layer neurons around GW33.

Inhibitory neuron expansion and migration into the developing neocortex occurs in parallel: Sst and Pax6 interneurons increased in number from GW10-25, Pvalb cells increased in number from GW20-30, while Lamp5 and Lamp5 Lhx6 cell types emerged at last in the third trimester. Gliogenesis occurred after the initial wave of neurogenesis, with OPCs and Astrocytes increasing in proportions from GW20 onwards and mature oligodendrocytes emerging only after birth (**Fig 1f, Supplementary Fig 2**).

In addition to changes in cell type proportions, we observed continuous transcriptomic maturation over age in each cell type, with increasing expression of corresponding marker gene sets in glutamatergic and GABAergic neurons (median spearman correlation between aggregate marker expression and age = 0.82, 0.78). We further examined whether newborn neurons had biased expression of specific subclass markers indicative of their future cell type (**Supplementary Figs 3-4)**. Re-annotating newborn neurons as mature subclasses based on relative marker enrichment revealed clear temporal patterns mirroring cortical neurogenesis: newborn glutamatergic neurons in late gestation were biased towards L2/3 IT (∼30-50% of cells at GW 33-38, **Supplementary Fig 3a-b**) while in early gestation, they were biased towards L6 IT (∼25% cells at GW15). Similarly, newborn GABAergic cells in late gestation were biased towards Pax6 (40-50% of cells at GW 33-37) while in early gestation they were biased towards Sst (30% of cells at GW15). Re-annotated newborn neurons also showed increasing expression of specific subclass markers with age (median R= 0.65) paralleling marker expression changes in mature subclasses (**Supplementary Fig 4**).

Of note, newborn neurons are immature and re-annotating them to mature subclasses resulted in highly heterogenous cell neighborhoods: LISI scores (local inverse Simpson’s Index^34^), a measure of heterogeneity of local cell neighborhoods in the UMAP were significantly higher for re-annotated newborn Glutamatergic and GABAergic neurons (mean LISI = 5.7, 4.4) compared to their original newborn identity (mean LISI = 1.77, 1.84, all pairwise Wilcoxon adjusted *P* < 10^−3^). In contrast, mature cell types had low LISI scores, indicating purity of their cell neighborhoods (mean LISI: 1.92 ± 0.1, **Supplementary Fig 3c**).

These results present an atlas of continuous cell type diversification and maturation in the developing human brain and serve as a starting point to further examine gene regulation underlying cell type maturation. To determine the trajectories with which different cell types approach their mature adult transcriptomic state, we next examined the replicability of cell types across datasets and timepoints.

### Replicable cell types show variable rates of maturation toward adult states

We used MetaNeighbor^35^ to assess the replicability of cell types across the human lifespan in an unbiased manner (**Fig 2a**). Cells of the same type were reliably grouped together across datasets and timepoints based on expression similarity (mean AUROC across all cell types= 0.78, **Supplementary table S2**). Vascular/Immune and Glial cell types showed highest replicability (mean = 0.96, 0.88), followed by Progenitors (mean = 0.8), mature GABAergic (mean = 0.8) and glutamatergic subclasses (mean = 0.74), and newborn Glutamatergic and GABAergic neurons (AUROC = 0.71, 0.78). The small number of unassigned cells in our dataset showed lowest replicability (0.53). Moderate replicability across cell types was observed in a more stringent test comparing each cell type to its next closest cell type (mean one vs next-best AUROC = 0.61).

**Figure 2:**
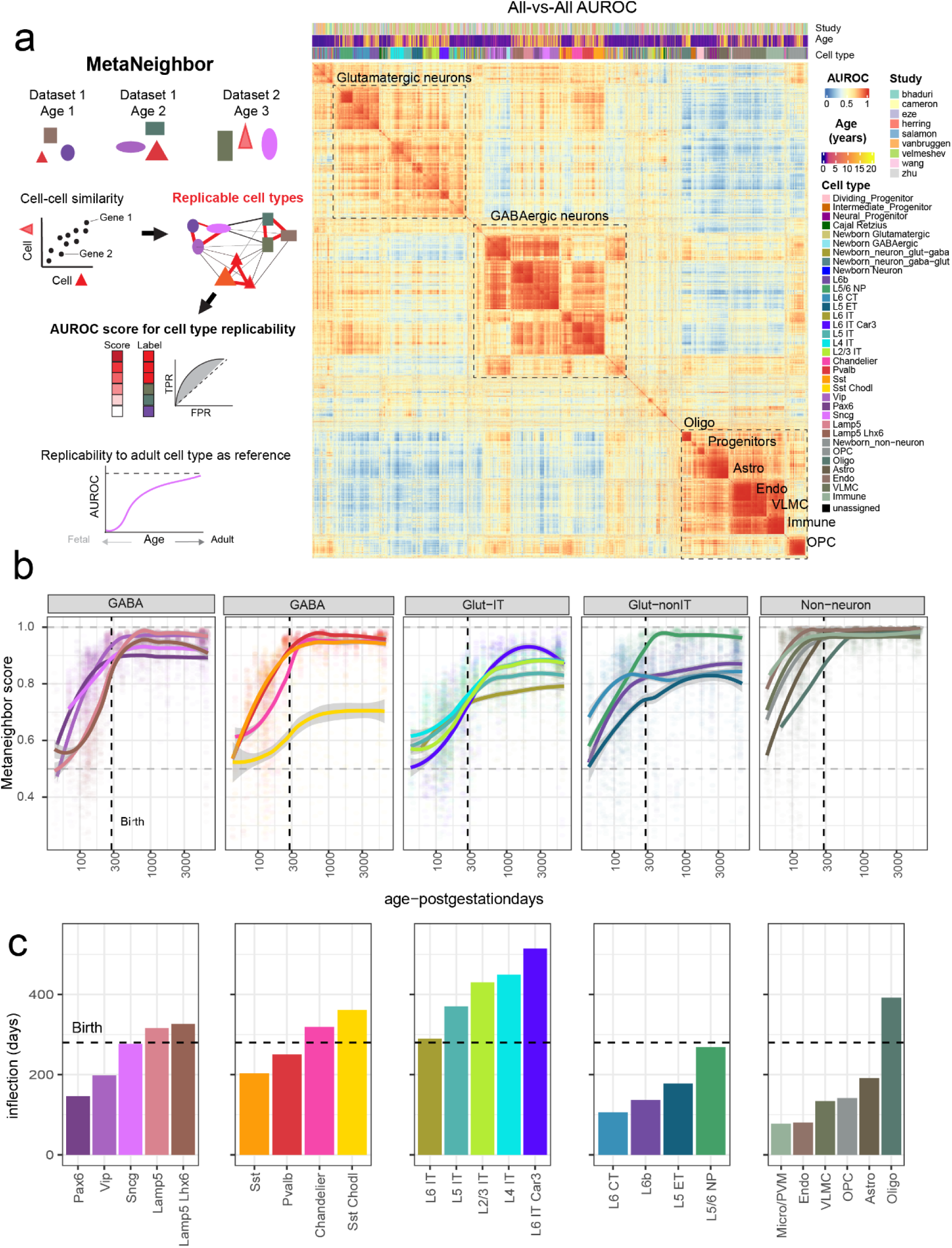
Cell types are replicable across timepoints, technologies, and studies and show variable rates of maturation towards adult state. **(a)** MetaNeighbor is used to assess cell type replicability across datasets and timepoints. AUROC scores measure the accuracy of predicting cell type for held out cells based on the cell-cell similarity network. Using the adult cell types as reference, we estimate the maturation rate of each cell type towards adult identity. (Right) All-vs-All MetaNeighbor AUROC heatmap clusters cells by cell type. **(b)** MetaNeighbor scores for each cell type at different timepoints with respect to the adult samples are plotted. **(c)** Inflection points denote the time at which fast phase of maturation towards adult state is completed for each cell type. Dotted lines denote birth.

To characterize the transition of cell types towards adult transcriptional states, we measured the replicability of cells from earlier developmental stages with respect to adult cells of the same type (**Fig 2b**). This evaluated whether the closest match for a cell type at a given timepoint is the adult version of that same cell type (score = 1) or whether it can be confused with other cell types (score = 0.5).

We identified clear cell type-specific temporal patterns with periods of rapid maturation that distinguished early and late cell types. Among glutamatergic neurons, L6 CT were the earliest maturing (**Fig 2c**), completing their first rapid phase of maturation by GW15, followed by L6b and L5 ET neurons which reached an inflection point around GW20-25. IT neurons (L6 IT, L6 IT Car3, L5 IT, L4 IT and L2/3 IT) were slower to mature and followed an inside-out pattern: L6 IT neurons reach an inflection point just around birth, while the other subtypes continued maturation postnatally, reaching their inflection points 3-8 months after birth. Among interneurons, Pax6 showed the earliest inflection (21 GW), followed by Vip (28 GW), while Sst and Pvalb cells continued rapid maturation until GW 29-35. Chandelier, Lamp5, and Lamp5 Lhx6 cells were the latest to mature, reaching their inflection 1-2 months after birth. Among non-neurons, vascular and immune cell types already have distinctive cell type-specific signatures by GW 11 which discriminate them from other cell types, while Oligodendrocytes were the latest to mature, reaching a stable state only 3-4 months after birth.

We further evaluated the replicability of cell types across the human lifespan to a recent dataset from the developing mouse visual cortex^17^ (**Supplementary Fig 5, Supplementary table S2**): all cell types were highly replicable between human and mouse for glutamatergic (9 subclasses, mean = 0.87) GABAergic (8 subclasses, excluding Pax6 which was not annotated in the mouse dataset^36^, mean = 0.94), glia (mean = 0.96), and vascular/immune cells (mean = 0.98). Comparing similar cell types also showed high replicability across species (mean one vs next-best AUROC = 0.69).

Comparing mouse cell type maturation with respect to adult human states revealed a faster pace of maturation: most mouse cell types were closely aligned to adult human states within 10 days after birth, with L2/3 IT neurons attaining a stable state by P17, while interneurons were transcriptomically stable earlier. These results are consistent with previous estimates of mouse corticogenesis^37^ and the known bradychrony of human neurodevelopment^38,39^.

Our results generally agree with the progression of inside-out cortical neurogenesis and gliogenesis and highlight temporal windows that are critical for cell type-specific maturation. Of note, lesser studied cell types including L5/6 NP and L6 IT Car3 were slow to mature in humans, which might be relevant in choosing the appropriate timepoint for their experimental investigation in other model systems.

### Spatial transcriptomic validation reveals emerging cortical layer identity of developing cell types

Next, to evaluate whether transcriptomic maturation is accompanied by refinement of cortical localization, we analyzed spatial transcriptomic datasets. Specifically, we used these datasets to register cell types annotated in our jointly processed atlas into age-matched MERFISH samples from the developing human cortex^26^ using Tangram^40^ (**Fig 3a**). 202,000 cells from GW14-16, 326,000 cells from GW20, and 63,000 cells from GW30-36 in the single-cell atlas were projected to GW15, GW20-22, and GW34 spatial samples respectively. This generated cell type probabilities for each cell in the MERFISH sample, with cells in the spatial dataset assigned to the best matching cell type from the single-cell atlas.

**Figure 3:**
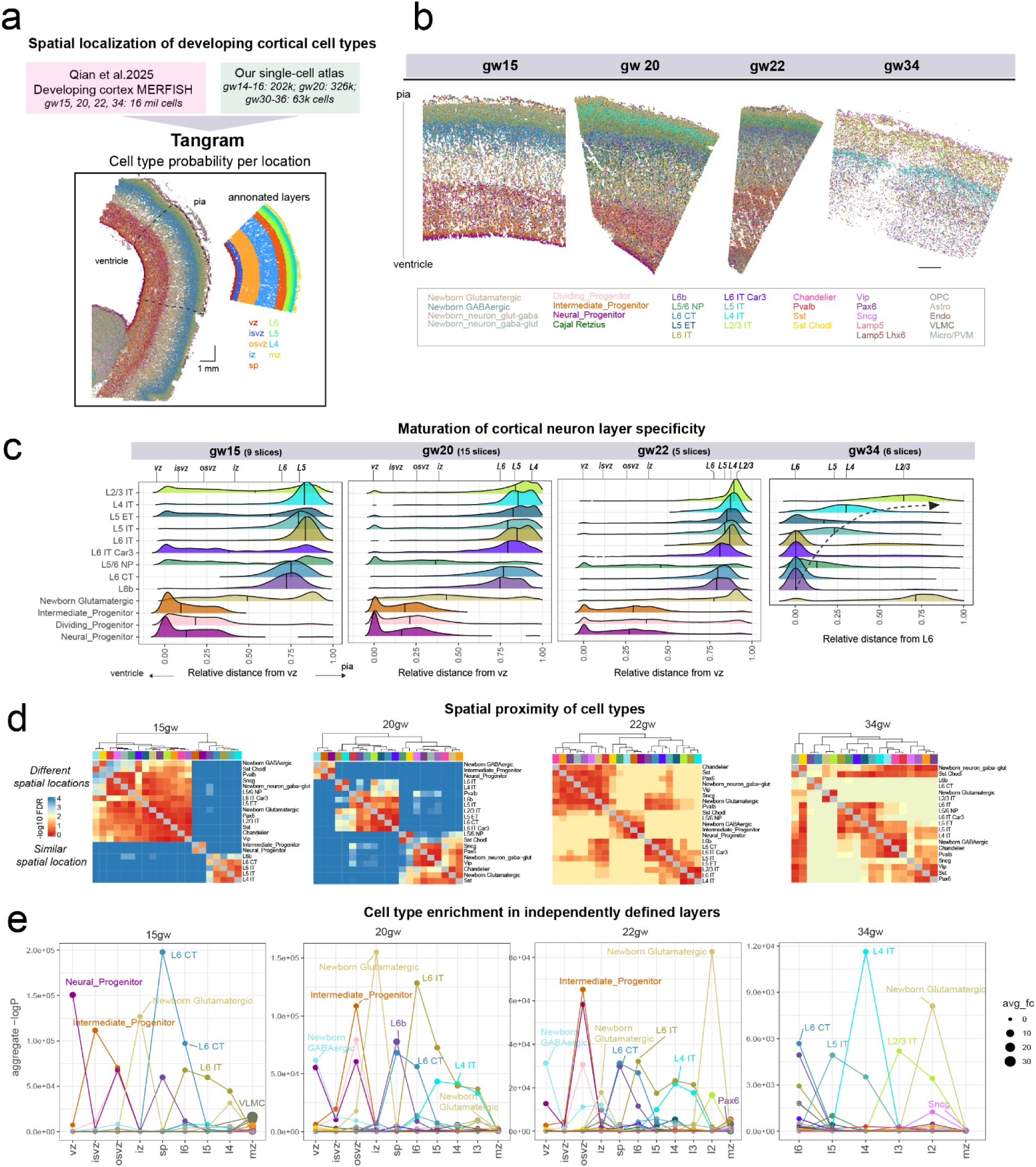
Spatial transcriptomic validation of cortical layer identity of developing cell types. **(a)** Our single-cell atlas is projected onto a MERFISH dataset^26^ from the developing human cortex at matched time points using Tangram^40^ to determine laminar organization of developing cell types. **(b)** Example MERFISH sections at different time points with cells colored by projected cell type annotations. **(c),** Ridgeplots show relative distance of excitatory neuron and progenitor cell types from the deepest ventricular zone. **(d)** Heatmaps show FDR adjusted p-values comparing relative cortical depth of each pair of cell types, blue: significantly different depths, red: similar depths. Older samples show higher resolution spatial clustering of cell types. **(e)** Aggregate P-values (y-axis) for enrichment of different cell types in independently defined cortical layers (x-axis). Cell types show improved laminar organization with age, with L2/3 identity emerging between 22-30 gestation weeks.

The laminar organization of cell types in the developing cortex became immediately evident: neural progenitors were dominantly present in the ventricular zone, newborn glutamatergic neurons in the intermediate zone, and L6 CT neurons in the emerging cortical plate at GW15; L4 IT cells appeared in the superficial layers at GW20-22 while superficial L2/3 IT cells were clearly visible only at GW34. Measuring the relative distances of each cell type from the deepest layer (ventricular zone or Layer 6), this analysis revealed clear inside-out migration of cell types with age (**Fig 3b-c**).

Pairwise comparisons of cortical depth between cell types also revealed emerging laminar organization over development (**Fig 3d**): at GW15 there were 3 broad clusters by spatial proximity: deep layer glutamatergic neurons, progenitors, and all other cell types; at GW20-22, newborn GABAergic neurons and neural progenitors were in the deepest ventricular zone, newborn glutamatergic neurons and GABAergic subtypes were within the intermediate zone, while mature glutamatergic subtypes were located more superficially in the cortical plate; finer-grained spatial domains were evident at GW34, with delineation of a L6b-L6CT zone, superficial L2/3 IT and newborn Glutamatergic zone, and subtype-specific interneuron laminar organization.

To further quantitatively assess whether our jointly processed cell types are spatially localized as expected, we relied on author-provided cortical zones (vz, isvz, osvz-ventricular, inner and outer subventricular zones, sp-subplate, l6-l2, mz-marginal zone) which were computed semi-manually independent of our cell type annotations. We computed enrichment of various cell types in each of these zones per slice and aggregated P-values across samples (**Fig 3e**). Using this supervised metric, we observed clear layer-specific enrichment. At GW15: Neural progenitors were in the vz, L6 CT, L6b cells were in sp, and L6 CT, L6 IT cells were in l5-6; at GW20: newborn GABAergic neurons were in vz, newborn glutamatergic neurons were in iz, and L4 IT in l4; at GW34: L2/3 IT and newborn glutamatergic neurons were in l2, and Sncg, Pax6 interneurons were in superficial marginal zones^41^.

These patterns reflect well established cortical laminar organization of cell types^23,42,43^ and independently validate our cell type annotation with a spatial reference. Given the clear maturation in cell type-specific laminar organization, we next sought to identify gene sets predictive of cell type-specific maturation.

### Cell type-specific maturation genes reveal developmental principles

To determine cell type-specific maturation gene programs within our atlas, we trained elastic net models that predict chronological age from each cell type’s transcriptome^27^. These transcriptomic ‘clocks’ narrow in on genes most predictive of age within each cell type, identifying coherent maturation programs (**Fig 4**). We successfully trained age-prediction models with high accuracy for most cell types in the prenatal brain (median correlation between predicted and actual age for 27 models = 0.78, median error = 3 weeks). Models use gene sets ranging in size from 13 (Lamp5) to 270 (L5 IT) per cell type with each gene having positive or negative coefficients (**Fig 4b**). Average expression of positive or negative maturation genes increase or decrease with age respectively (**Fig 4d-e**).

**Figure 4:**
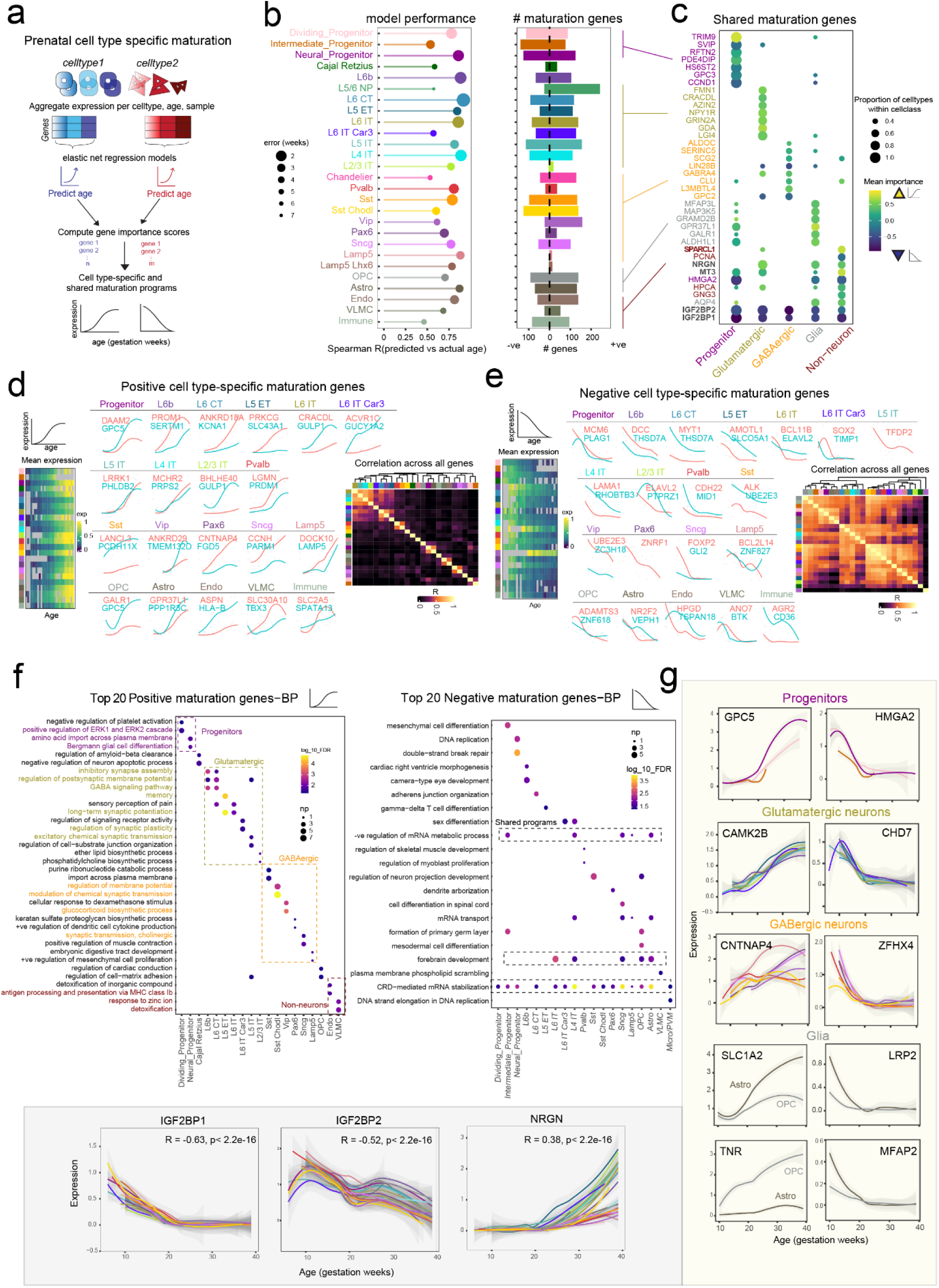
Cell type-specific transcriptional maturation programs in the developing human brain. **(a)** Framework for identifying cell type-specific maturation programs. Gene expression is aggregated by cell type, age, and sample and cell type-specific elastic net regression models are trained to predict age from gene expression. Gene importance scores are derived for each model (bootstrap measure, see Methods). These scores identify top cell type-specific maturation programs. **(b)** *(Left)* Model performance for each cell type in the fetal brain, quantified as Spearman’s R between predicted and actual gestational age. Dot size reflects the prediction error in weeks. *(Right)* Number of positive- and negative-weighted maturation genes per cell type. **(c)** Shared maturation genes in major cell classes. Each dot represents a gene, with dot size indicating the proportion of cell types in which the gene is identified as a maturation gene, and color indicating the mean importance score in that cell class (yellow = positive/upregulated; blue = negative/downregulated). **(d)** Heatmap *(left)* displays mean expression levels of all positively weighted genes per cell type across age. *(middle)* Top cell type-specific genes’ expression trajectory are shown for each cell type. Correlation matrix (*right*) shows pairwise Spearman correlations of all gene importance scores across cell types. **(e)** Layout as in (d), but for negatively weighted genes. **(f)** Gene Ontology (GO) enrichment analysis for the top 20 positive *(left)* and negative *(right)* maturation gene sets across cell types. Dot color represents –log_10_(FDR), and size indicates number of genes intersecting that GO term. Highlighted terms indicate representative processes per class. **(g)** Representative maturation gene trajectories: Expression is plotted against gestational age (weeks) for Progenitors (*GPC5*, *HMGA2*), Glutamatergic neurons (*CAMK2B*, *CHD7*), GABAergic neurons (*CNTNAP4*, *ZFHX4*), Astrocytes, and OPCs (*SLC1A2*, *LRP2*, *TNR*, and *MFAP2)*. Shaded ribbons are 95% confidence intervals. *(Bottom)* Expression trajectories for shared maturation genes: *IGF2BP1*, *IGF2BP2*, and *NRGN* across all cell types.

We assessed the importance of maturation genes in each cell type-specific model by evaluating gene usage frequency across bootstraps (**Supplementary table S3**). Genes with high bootstrap importance scores tend to be most highly correlated to age in that cell type (**Supplementary Fig 6-7**). Maturation programs within each cell class are highlighted in **Fig 4c** and include well-known synaptic receptors and dendritic molecules (*GRIN2A*, *NPY1R*, *GABRA4*, *NRGN*, *GDA, CLU*^44–49^) in glutamatergic and GABAergic neurons, canonical glial (*AQP4, ALDHL1*^50,51^), vascular (*SPARCL1, MT3*), and neural progenitor markers (*CCND1*^52^*, HMGA2*^53^). *IGF2BP1* and *IGF2BP2* were globally downregulated across all cell types and known to be critical for neurodevelopmental timing^54,55^ (**Fig 4c**).

We next identified cell type-specific maturation genes by prioritizing genes with high bootstrap importance whose expression is specifically correlated to age only within that cell type (see Methods, **Supplementary Fig 7, Fig 4d-e, Supplementary table S4**). Developmentally upregulated genes (i.e. with positive coefficients) include markers and biological processes relevant to that cell type: e.g. *CPLX3, PROM1* in L6b neurons^56,57^, ion channel *KCNA1* in L6 CT neurons, *PRKCG* (*protein kinase C gamma*) - a corticospinal tract marker in L5 ET neurons, *GULP1-* an amyloid precursor protein processing molecule^58^ in IT neuronal classes, upper-layer neuron-specific TF (transcription factor) *BHLHE40*^59^ in L2/3 IT neurons, median ganglionic eminence TF *PRDM1* in Pvalb cells^3^, *PCDH11X* in Sst, *ANKRD29* in Vip, neurexin *CNTNAP4*^60^ in Pax6, *CHRNA2* (nicotinic acetylcholine receptor) and marker gene *PARM1*^61,62^ in Sncg, and *LAMP5* in Lamp5 interneurons. Among non-neurons, these include *PPP1R3C* (or *PTG*, protein targeting to glycogen) - a regulator of glycogen metabolism in astrocytes^63^, extracellular matrix protein-encoding *FBLN1 (Fibulin1)* and *GALR1* (*galanin receptor 1*) which regulate OPC maturation^64,65^, *HLA-B* in endothelial cells, and the microglia-specific fructose transporter gene *SLC2A5*^66,67^.

Positive maturation genes in 17/24 cell types were significantly enriched for that cell type’s markers. Gene importance scores across all positive maturation genes (**Supplementary table S5**) show poor correlation between cell types, highlighting the specificity of this gene set (**Fig 4d, inset**). Enriched biological processes include relevant cell type-specific programs: positive regulation of ERK1-2 cascade, amino acid transport, and glial cell differentiation in progenitors indicating a progressive switch towards gliogenesis; neuronal terms including synapse assembly, synaptic plasticity, regulation of membrane potential in glutamatergic and GABAergic neurons, and antigen presenting and detoxification in Endo and VLMC cells respectively (**Fig 4f**).

In contrast, developmentally downregulated genes (**Fig 4e, Supplementary Fig 8**) represent general cellular processes including early stem cell and proliferative markers such as *MCM6* (DNA replication), *PLAG1*, *SHISA2* (Wnt inhibitor) in progenitors, *ADAMTS3*^68^ (promoter of OPC proliferation) in OPCs, neural stem cell marker *VEPH1*^69^ in astrocytes, neural progenitor markers *SOX2*, *PTPRZ1* and the subcortical projection-specifying TF *BCL11B*^70^ in IT neurons, receptor for axon guidance molecule Netrin-1 (*DCC*) in L6b, L6 CT and L5 ET neurons, protein degradation pathway genes - *UBE2E3, MID1, ZNRF1* in Sst, Vip, Pvalb, and Pax6 cells, TF *FOXP2* in Sncg, DNA-damage response gene *ZNF827* in Lamp5, angiogenesis regulator *TSPAN18* in Endo, and *CD36* in Immune cells. Overall, negative maturation genes show zero overlap with cell type-specific marker genes and are highly correlated in their gene importance scores across cell types (**Fig 4e**). Enriched biological processes include shared programs such as negative regulation of mRNA metabolism and translation, mRNA stabilization, in addition to stem cell population maintenance, proliferation, and differentiation programs across cell types (**Fig 4f**).

### Genes important for cell type maturation are evolutionarily constrained

Consistent with their more general program, negative maturation genes are more ubiquitously expressed across cell types in first trimester (Wilcoxon p-value < 2.2 *10^−16^) and are evolutionarily constrained: As gene importance increases, negative maturation genes show greater evolutionary constraint, with top 25% showing significantly lower LOEUF scores^71,72^ (mean = 0.66) compared to positive genes (mean = 0.79). In contrast, positive maturation genes show increasing overlap with SFARI genes as their gene importance increases: top 25% show significantly greater overlap with SFARI genes compared to negative genes (24% vs 8% mean overlap, **Supplementary Fig 9**). We further evaluated the overlap with autism-linked genes with rare variants: SFARI score 1, as well as 185 ASD genes and 664 NDD (neurodevelopmental disorder) genes linked to developmental delay from Fu et al. 2022^73^. Positive maturation genes showed greater overlap with SFARI score 1 and ASD185 gene sets compared to negative genes, but both positive and negative maturation genes showed equivalent and higher overlap with the broader NDD664 gene set (**Supplementary Fig 9b, Supplementary table S6**). For example, negative maturation genes in L6 IT, L5 IT cells and positive maturation genes in L6 IT Car3, L5 ET cells were significantly enriched for NDD644 genes. These results indicate distinct expression trends of neurodevelopmental disorder risk genes and reveal higher evolutionary constraint of negative maturation genes.

Cell type-specific maturation programs further highlight the biological transition from stem cells and progenitors to differentiated cell types. For example, in neural progenitors the proliferative marker *HMGA2* decreases^15^ while glial marker *GPC5*^74^ increases, reflecting the transition from neural proliferation to gliogenesis. Glutamatergic neurons show increasing expression of excitatory signaling molecule *CAMK2B*, and decreasing expression of *CHD7*^75^, a chromatin remodeler and regulator of neuronal differentiation. Similarly, in GABAergic neurons, neurexin *CNTNAP4*^76^, critical for GABA release, increases in expression while the TF *ZFHX4* which regulates differentiation decreases. Astrocytes show increasing expression of the glutamate reuptake transporter *SLC1A2* while OPCs show increasing *TNR* which promotes OPC differentiation to Oligodendrocytes^77^. In addition, conserved m6A-mRNA reader molecules *IGF2BP1, IGF2BP2*^54,55^ are downregulated uniformly across all cell types (**Fig 4g**).

In summary, our cell type-specific transcriptomic clocks and gene importance scoring approach identify molecular drivers of cell-type maturation, revealing differences in evolutionary conservation and disease enrichment between up- and down-regulated genes. All gene importance scores and gene expression trajectories within cell types can be explored in our <u>brain-development browser</u>.

### Evaluation of immune cell trajectories in the developing brain

As a further application of our atlas, we next examined the immune cell population, annotating its diversity in greater detail and constructing pseudotime trajectories to evaluate maturation within these rare cell types. The earliest microglia in the human embryonic brain appear at GW4 from progenitors in the yolk sac^78,79^ and complete their colonization by week 13, when the blood brain barrier is fully formed. They play a vital role in surveillance, synaptic support and pruning, innate immune defense, myelination and phagocytosis, and have specialized functions at various stages of development.

While the diversity of microglia in activated and disease conditions has been extensively studied, their developmental trajectories in the human brain remain incompletely characterized.

We subclustered 35,111 cells identified as Immune cells into three major cell types: microglia, macrophages and lymphocytes (**Fig 5a-d**). Within the microglia, many clusters corresponded to stages of development and could be broadly classified into nascent, intermediate, and mature populations.

**Figure 5:**
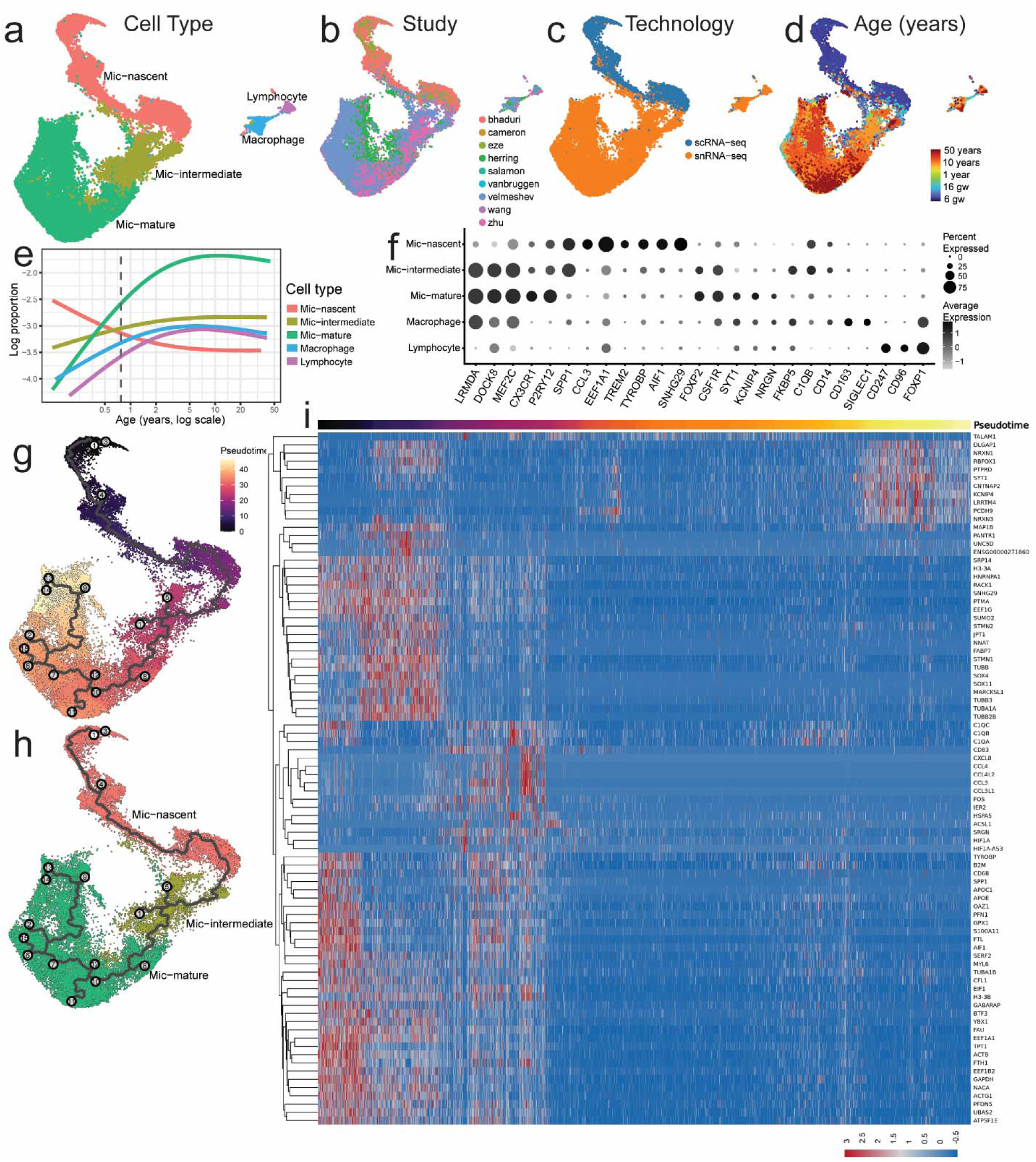
Exploring immune cell diversity in the developing human brain. **(a)** Populations of microglia (Mic), macrophage and lymphocytes seen in the human brain across **(b)** Studies, **(c)**Technology and **(d)** Age. **(e)** Log of the proportions of the cells in each subgroup with respect to the total number of cells from each donor across time. Vertical dashed line denotes birth. **(f)** Dotplot displaying top marker genes for each subgroup. **(g-h)** Monocle based trajectory displaying the branching in the broad microglial states across pseudotime. **(i)** Branched heatmap of genes driving the trajectory of microglial maturation against pseudotime.

**Figure 6:**
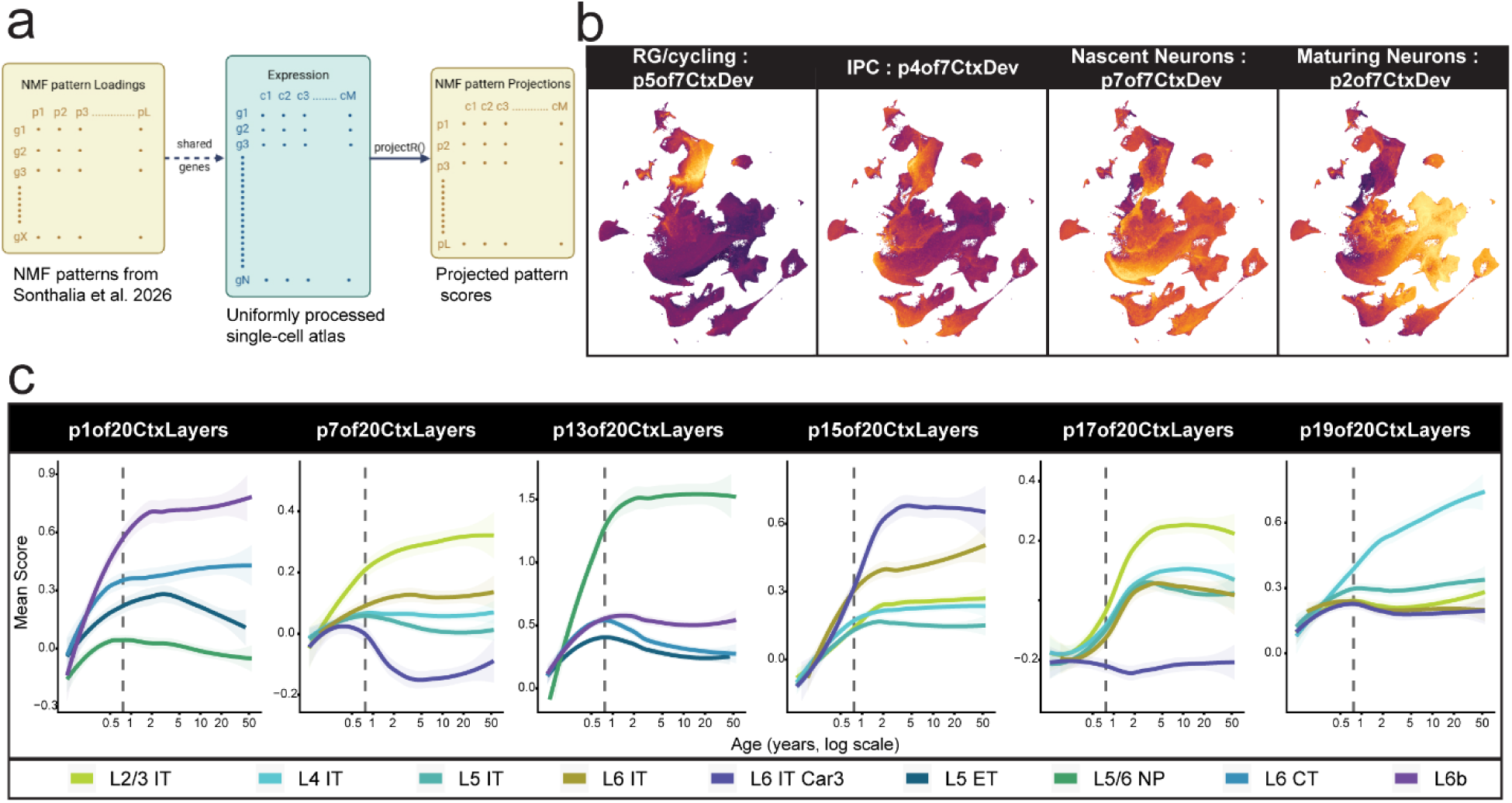
Validation of gene modules corresponding to early neurogenesis and cortical layer identity. **(a)** NMF pattern projection strategy. Pattern loadings are obtained from Sonthalia et. al. (2026) for the p7 and p40 cortical developmental data and the p20 cortical layers data. Shared genes are identified with the expression matrices of our atlas and NMF patterns are projected using the projectR package in R. **(b)** UMAPs colored by the projection scores of patterns corresponding to radial glia (RG), intermediate progenitor cells (IPC), nascent and maturing neurons. **(c)** Scores of cortical layer cells for highlighted NMF patterns across age. Vertical dashed line denotes birth.

Through a generalized additive model, we determined the association between the proportion of microglial cell states and the age of the donors. As expected, we found a downward trend for nascent microglia and an upward trend for mature microglia (**Fig 5e**). The mature signatures were finally attained during mid-childhood, but ∼10% of microglia from adult donors remained in the intermediate states.

Mic-nascent were characterized by high expression of genes encoding ribosomal proteins, structural proteins (*TUBB, TUBB2B, ACTG1*), and transcriptional machinery (*EEF1A1, EEF1G, EIF1*), as well as low levels of microglia specific genes (*AIF1, P2RY12, SPP1*) (**Fig 5f, Supplementary table S7**). Since microglia are derived from yolk sac hematopoietic progenitors, we compared microglial states in the brain to cell types identified in a human yolk sac atlas at GW4-8^79^. MetaNeighbor analyses revealed the highest replicability of Mic-nascent with Common Myeloid Progenitor (CMP) populations (AUROC = 0.97) and Hematopoietic Stem and Progenitor Cells (HSPC) (AUROC = 0.92). Thus, Mic-nascent represents a relatively homogeneous population of nascent microglia that retain transcriptional signatures of yolk sac-derived progenitors.

Mic-intermediate represented a more diverse group of immature microglia that could be sub-clustered into 15 distinct groups, all of which expressed canonical microglial markers at varying levels. These populations were distinguished by the expression of markers associated with early proliferative states (MicImmature - 1, 4, 10), dividing cells (MicImmature-14), axon tract microglia (MicImmature - 3, 5, 7), reactive cells (MicImmature-11), and a monocyte-like population (MicImmature - 6) (**Supplementary Fig 10 a, b, Supplementary table S8**).

Sub-clustering of Mic-mature cells revealed 12 groups. The marker genes for these groups overlap established signatures of homeostatic (Mic-mature - 0, 1, 2, 5), Disease Associated (Mic-mature - 6), activated (Mic-mature - 7), phagocytic (Mic-mature - 11) and interferon responsive microglial states (Mic-mature - 12). An intriguing population that was most abundant during early childhood expressed a gradient of synapse-specific genes, potentially contributing to the interactions of microglia with synapses (Mic-mature - 3, 4, 8, 10), (**Supplementary Fig 10 c, d, Supplementary table S9**).

We constructed a continuous trajectory to characterize the sequence of maturational states, rooted in Mic-nascent. The trajectory extended through Mic-intermediate, then through multiple branches leading to distinct states of mature microglia (**Fig 5g-h**). Genes enriched at the beginning of this trajectory included ribosomal components, cytoskeletal genes and genes involved in protein synthesis (**Fig 5i, Supplementary table S10**). This was followed by cell surface proteins such as *UNC5D*, which has a known function in axonal guidance^80^ and embryonic transcription factors *SOX4* and *SOX11*. Several canonical microglial markers, including *LRMDA* and *DOCK8,* were expressed only at postnatal cells at higher pseudotime, representing signatures that distinguish mature populations from nascent and intermediate states. We also identify *PCDH9-*expressing mature microglia at later pseudotimes, consistent with previous reports^81^. Overall, these analyses reveal the transcriptional states of microglia throughout development and demonstrate the dynamic quality of immune signatures evident in our jointly processed atlas.

### Validation of cell type and layer-specific gene programs

Going beyond individual cell type and gene-centric approaches, we next examined whether transcriptome-wide signatures of specific cell types and cortical layers^28^ could be recapitulated in our atlas.

We projected cortical development and cortical layer NMF pattern loadings from Sonthalia et. al., 2026^28^ into the expression data of all 2.16 million cells. Several of the highlighted biologically meaningful patterns seen in their publication corresponded to cell types within our integrative dataset. We used these projections to determine temporal dynamics of cell type- and layer-specific patterns through development.

Among cortical development patterns, P5of7 was the radial glia/cycling pattern which was highest expressed in a subset of the neural and dividing progenitors. P4of7 corresponding to IPCs was expressed in the intermediate progenitor cells, P7of7 for nascent neurons had highest expression in the newborn glutamatergic cells and P2of7 specific to maturing neurons was highly expressed in all the cortical layer cell types (L2/3 IT, L4 IT, L5 IT, L6 IT, L6 IT Car3, L5 ET, L6 CT, L6b, L5/6 NP).

They also described 20 cortical layer patterns derived from adult cortical datasets, of which several corresponded to individual cortical layer assignments in our dataset. P1of20 peaks in L6b in early infancy; P7of20 in L2/3 IT gradually increases through prenatal development, while p17of20 is switched on during infancy in L2/3 IT, L4 IT, L5 IT and L6 IT. P13of20 is switched on at birth in L5/6 NP; P15of20 is L6 IT specific, particularly L6 IT Car3 starting during early childhood and P19of20 gradually increases in L4 IT.

These results indicate that transcriptome-wide programs of neural progenitor, maturing neuron, and cortical layer identity show consistent expression in our atlas following the timeline of cell type maturation described in **Fig 2**. Having established that known developmental programs are recoverable, we next explored how gene regulatory dynamics are coordinated across cell types in the developing brain.

### Multi-scale gene regulation in the developing brain

Gene regulation in the developing brain directs cell fate specification as well as maturation within individual cell types. A key question is the scale at which these gene regulatory programs operate (**Fig 7a**): co-expression may be shared across multiple scales (coherent both across and within cell types), restricted to between-cell-type differences (across-cell types only), or specific to regulation within individual cell types (within-cell types only)^82^. Distinct co-expression patterns across vs within-cell types would indicate rewiring of gene regulatory circuits at different scales whereas coherent multi-scale gene regulation would imply differential usage of the same core gene-regulatory program at different scales.

**Figure 7:**
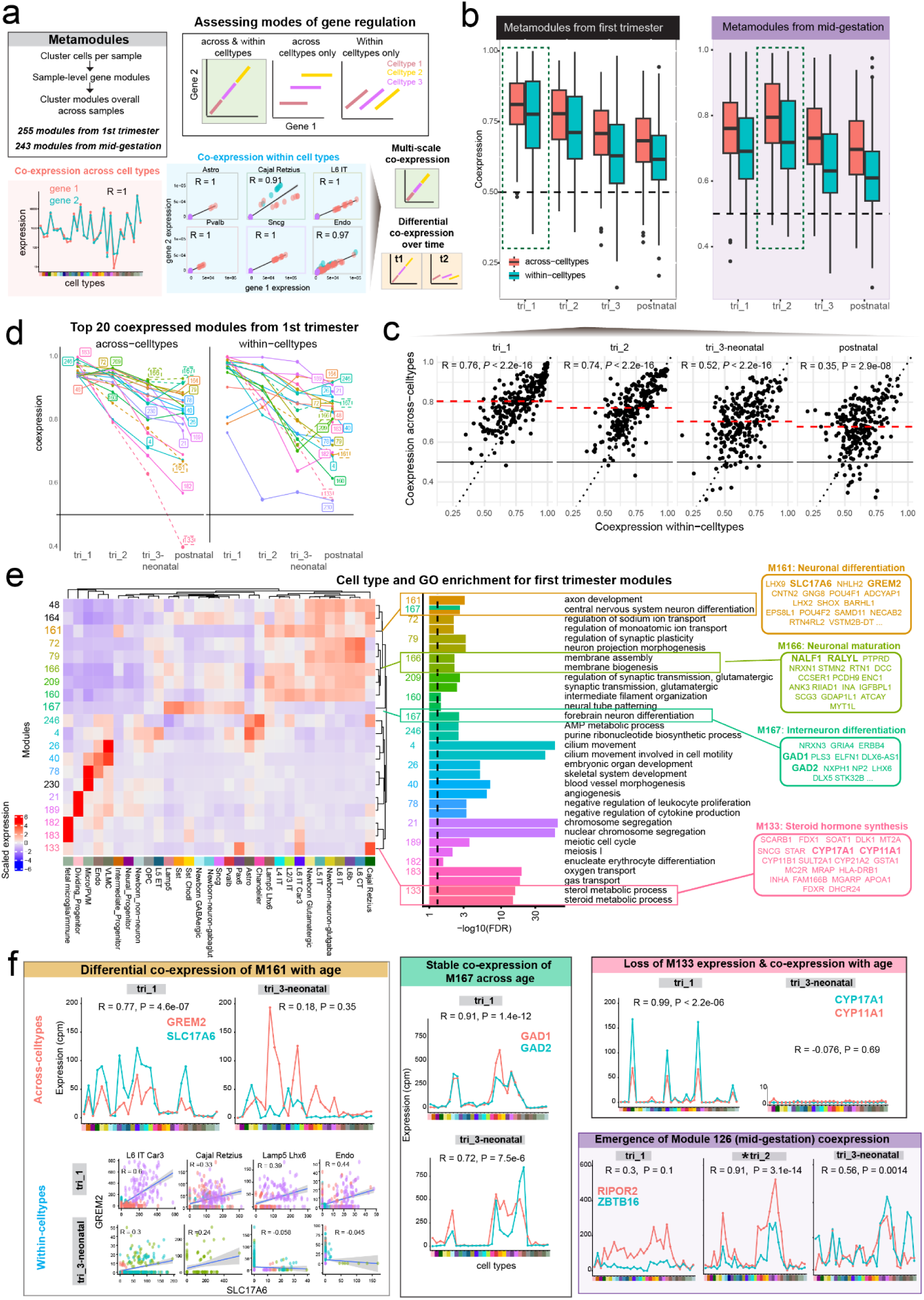
Multi-scale co-expression analysis reveals cell type-specific and temporally dynamic gene regulatory programs in the human brain. **(a)** Metamodule generation^29^ and co-expression analysis strategy. *(Top)* Three modes of gene co-expression are distinguished: across and within-cell types, across-cell types only, and within-cell types only, illustrated by divergent slopes of gene pair co-expression across cell type contexts. *(Bottom)* Example of co-expression across cell types, where two genes show correlated expression (R = 1) uniformly across all cell types (*left*) and example of within-cell-type co-expression, where the gene pair is also correlated within individual cell types (*middle*). This framework lets us evaluate multi-scale co-expression and differential co-expression over time. **(b)** Box plots comparing across-cell-type (red) and within-cell-type (blue) co-expression scores for metamodules derived from first trimester *(left)* and mid-gestation *(right)* across four developmental stages. **(c)** Scatter plots of across vs. within-cell-type co-expression for first-trimester metamodules. Dashed red lines indicate mean across-celltype co-expression. **(d)** Differential co-expression of top 20 metamodules from the first trimester across developmental stages. **(e)** Cell type and Gene Ontology (GO) enrichment for first-trimester metamodules. Heatmap shows expression of selected metamodules across cell types. Bar plots of the top enriched GO terms for each module and highlighted genes from four modules: M161 (neuronal differentiation), M166 (neuronal maturation), M167 (interneuron differentiation), and M133 (steroid hormone synthesis) are shown to the right. **(f)** Examples illustrate distinct co-expression dynamics: *(Left)* Module M161 genes *GREM2* and *SLC17A6* are co-expressed at tri_1 but not at tri_3–neonatal across and within cell types. *(Center)* Module M167 genes *GAD1* and *GAD2* maintain stable co-expression over time at both tri_1 and tri_3–neonatal. *(Top right)* Loss of M133 expression and co-expression with age: *CYP17A1* and *CYP11A1* are co-expressed at tri_1 but not at tri_3–neonatal, due to loss of expression. *(Bottom right)* Emergence of Module 126 (from mid-gestation) co-expression: *RIPOR2* and *ZBTB16* show negligible co-expression at tri_1 but strong co-expression emerging at tri_2 and maintained at tri_3–neonatal.

We explored this by deriving developmental meta-modules^29^ from first trimester (< 14GW, 255 modules each with 10-436 genes, **Supplementary table S11**) and mid-gestation (20–25 GW, 243 modules each with 10-551 genes, **Supplementary table S12**). Each module represents genes that are co-regulated across all the individuals in our atlas. We then evaluated module co-expression strength within and across cell types in aggregate co-expression networks at specific developmental stages using EGAD^83^ (see Methods). Across both sets of gene modules, co-expression strength across and within cell types was highly correlated (**Fig 7b-c**) with extensive temporal remodeling: first trimester modules were highly co-expressed across and within-cell types in the first trimester (mean AUROC = 0.81, 0.77, R = 0.76, *P* < 2.2 * 10^−16^), but showed substantially lower co-expression in postnatal periods (mean AUROC = 0.68, 0.62, R = 0.35, *P* = 2.9 * 10^−08^). Similarly, modules derived from mid-gestation showed highest across and within-cell type co-expression in second trimester (mean AUROC = 0.80, 0.73, R = 0.8, *P* < 2.2 * 10^−16^), but weaker co-expression in third trimester-neonatal stages (mean AUROC = 0.73, 0.65, R = 0.7, *P <* 2.2 * 10^−16^, **Fig 10**). These results support temporally dynamic multi-scale gene regulation in the developing human brain.

Focusing on the top 20 highest co-expressed modules in the first trimester (**Fig 7d**), we identified both cell type-specific and broadly expressed modules enriched for distinct biological processes (**Fig 7e**): Module 21 and 189, expressed in Dividing Progenitors are enriched for cell cycle terms; Module 4 expressed in Astrocytes, Chandelier cells, and L5 ET is enriched for cilium movement; Module 133 expressed in Cajal Retzius and a few other cell types is composed of enzymes from the steroid hormone synthesis pathway (e.g. *CYP17A1*, *CYP11A1*); Module 161, broadly expressed across glutamatergic cell types is enriched for neuronal differentiation (e.g. vesicular glutamate transporter *SLC17A6*, and BMP antagonist *GREM2*^84^) while Module 166 is enriched for neuronal maturation-related genes (e.g. *NRXN1*, *PCDH9*); Module 167, expressed across GABAergic cell types is enriched for forebrain neuron differentiation and includes several lineage markers (e.g. *GAD1, GAD2, DLX5*).

Several modules exhibited differential co-expression over development (**Fig 7e**): Neuronal differentiation module 161 is highly co-expressed in first trimester, with genes *SLC17A6* and *GREM2* showing correlated expression across and within cell types (**Fig 7f, left**), but this relationship is disrupted by the third trimester-neonatal stage despite both genes being expressed at comparable levels. Steroid hormone synthesis module 133 is also tightly co-expressed in the first trimester, but the expression of genes *CYP17A1* and *CYP11A1* falls to near-zero by the third trimester, eliminating their co-expression (**Fig 7f, right**). Intriguingly, module 133 is expressed in Cajal-retzius cells and the early co-expression of the steroidogenesis pathway in this population may have functional implications for corticogenesis^107,108^. Likewise, a mid-gestation module expressed in CGE-interneurons is not co-expressed in the first trimester, with genes *ZBTB16* and *RIPOR2* becoming co-expressed only from the second trimester onward (example mid-gestation modules are highlighted in **Supplementary Fig 11**). In contrast, interneuron differentiation module 167 (*GAD1, GAD2*) shows stable co-expression across all developmental stages (**Fig 7f, middle**).

Together, these results demonstrate that our framework recovers gene regulatory relationships that are consistent within and across cell types, and delineates how co-expression is gained, maintained, or lost over the course of brain development.

From early observations of correlated ion channel gene expression across neuronal cell types^85^, to the convergence of co-expression signatures in neurodevelopmental disorders^86–88^, the identification of co-expression patterns that predict cell state, disease, and developmental transitions remains a central goal in developmental neuroscience. Here, we define cell type-resolved and temporally dynamic gene modules that mark fundamental biological transitions and offer tractable targets for future experiments. The full set of developmental modules is available to explore through our <u>brain-development</u> <u>browser</u>, providing a community resource to guide prioritization of experimental targets.

## Discussion

Brain cell atlasing efforts have catalogued human neural cell type diversity across different developmental stages in great detail^1–3,6,9,14,23,89^. However, identifying consensus biological principles from these independent datasets to pursue the next generation of mechanistic questions in developmental neuroscience has remained difficult. Here, we present a consolidated atlas of human brain development from GW6 to adulthood, reprocessed from raw sequencing reads and annotated with a principled cell-type framework spanning 2.16 million cells, 156 donors, and nine studies. Our analysis yields highly replicable cell-types across the lifespan, enabling us to move beyond atlas-building to identify critical periods in cellular maturation and the dynamic gene programs that shape cell fate and cell type–specific trajectories.

While human neurodevelopment is widely acknowledged to be a continuous and slow process^90^, the timing at which individual cell types acquire adult-like transcriptional states remains unresolved, including whether this transition occurs before or after birth. Atlases sampling distinct developmental stages have arrived at substantially different estimates: Bhaduri et al. found poor concordance between second-trimester and adult transcriptional states^1,15^, whereas Braun et al. detected signatures of adult midbrain neuron identity in first trimester samples as early as GW10^14^; Herring et al. profiled the prefrontal cortex from GW22 onward and reported that all cell types transitioned towards adult states only postnatally with deep-layer neurons maturing in infancy and upper-layer neurons maturing in adolescence^6^. Velmeshev et al. used pseudotime to quantify maturation from second trimester to adulthood and found two broad groups: L5, L5/6 IT, and interneurons were largely mature by second trimester, while L2/3, L4 IT, and L6 neurons continued to mature postnatally^2^.

We quantified the transition to adult states across cell types on a common temporal axis by leveraging our jointly processed atlas: Assessing each cell type’s transcriptomic similarity to its adult counterpart across studies, we found that non-neuronal cells (Endo, VLMC, Immune-Micro/PVM), deep-layer glutamatergic neurons (L5-6), and specific interneurons (Pax6, Vip, Sst, Pvalb) approached adult-like states prenatally, while upper-layer IT neurons, Lamp5, Chandelier interneurons, and oligodendrocytes matured postnatally within the first year after birth. Intriguingly, lesser-studied L6 IT Car3 and L5/6 NP cells were among the slowest-maturing glutamatergic cell types: L6 IT Car3 neurons share transcriptomic identity and long-range cortico-cortical projection patterns with claustrum neurons^91,92^, while L5/6 NP neurons form a rare locally-projecting subclass^92–94^, with a broad repertoire of neuromodulatory receptors^57^ and are selectively responsive to psilocybin exposure in mice^95^. Their protracted maturation likely depends on interactions with surrounding cortical circuitry and warrants further functional investigation. Furthermore, comparing human cell type maturation against the mouse brain^17^ revealed accelerated maturation across cell types in mice, consistent with the prolonged timeline of human neurodevelopment ^39,96^.

Understanding the specification and expansion of immune cell lineages in the human brain is of increasing interest given their role in normal neurodevelopment^97^. While prior atlases have sampled microglia from limited prenatal time points^98–100^ and adult or diseased samples^101^, our atlas recovers over 35,000 immune cells across all stages of the developing human brain. This enabled us to resolve heterogenous nascent, intermediate, and mature microglial states, including rare sub-groups. Our analyses resolve the timing of these developmental transcriptional states in unprecedented detail.

Our cell type-specific transcriptomic clocks and co-expression analysis identify genes with continuous expression trends critical for cell type maturation, and gene-modules that switch on/off at sensitive time points. Several of the cell type-specific maturation genes we identified including *CLU*^48,102^*, GABRA4*^103^, and *ZFHX4*^104^ are implicated in neurodevelopmental disorders. Beyond canonical marker genes, ion channels, synaptic receptors, and axon guidance molecules (*PLXND1*^105^), we find *GULP1*, an amyloid precursor protein adaptor^58^ to be developmentally upregulated across glutamatergic IT neurons, and *SRGAP2B*, a human-specific paralog previously shown to drive microglial morphological complexity^106^, to be upregulated in microglia. Further insights relevant to specific cell types can be identified by exploring our maturation gene sets in detail.

More broadly, our analysis reveals two complementary developmental principles: developmentally upregulated genes are cell-type-specific and strongly enriched for autism risk genes, whereas developmentally downregulated genes are proliferation and differentiation-associated, shared across cell types, and under stronger evolutionary sequence constraint. Overall, cell type-specific maturation genes were enriched for broad neurodevelopmental disorder risk genes. At the module level, we observed differential co-expression of programs underlying neuronal differentiation and maturation, as well as striking first trimester-specific co-expression of the steroid hormone synthesis pathway in Cajal-Retzius neurons. While steroidogenesis has been demonstrated in preplate Cajal-Retzius and subplate neurons in mice and specific neuronal populations in rats^107,108^, its role in early human brain development is largely underexplored and our analysis provides a starting point for future experimental investigation of this pathway.

As a summarizing observation, projecting our jointly processed cell-type annotations onto developmental spatial transcriptomic data revealed clear changes in laminar localization that closely recapitulate the well-established inside-out pattern of cortical development. This finding is somewhat unexpected, as our prenatal subclass annotations are derived from marker genes identified at adult time points. The observation that prenatal cells annotated using adult markers faithfully capture known developmental layer dynamics strongly suggests that core features of adult molecular identity are established remarkably early following neuronal differentiation. Together with the protracted, continuous expression changes we observe across the lifespan, these results support a model in which neurons rapidly acquire their defining cell-type transcriptional programs early in development, followed by gradual refinement and maturation over time. Notably, the extent and trajectory of this later refinement appear to vary across cell types.

Several caveats apply: We did not analyze inter-regional differences in detail here because cross-study comparisons were underpowered, with any given region sampled in at most two studies. We also did not resolve higher-resolution cluster or subtype identities, focusing instead on pre-defined subclasses that are highly replicable across datasets and conserved between human and mouse.

By harmonizing nine independent datasets into a continuous atlas of cell type diversity in the human brain across the lifespan, we establish a foundational reference for future comparisons across *in vitro* models, species, and disease states. The accompanying <u>brain-development browser</u> with maturation gene modules and co-expression explorer will serve as a valuable hypothesis-generating resource for future experimental investigations.

## Methods

### Data processing

FASTQ files for all 9 studies were obtained from NEMO Archive (https://nemoarchive.org/) or directly from the authors and processed using the Optimus pipeline^109^ (RRID:SCR_018908, https://broadinstitute.github.io/warp/docs/Pipelines/Optimus_Pipeline/README) developed for the BRAIN Initiative Cell AtlasNetwork (BICAN). Each sample was filtered to include cells with a minimum of 500 genes.To avoid removing maturing cell types with elevated mitochondrial content, prenatal cells were not filtered by mitochondrial gene percentage; the distribution of mitochondrial gene percentages across studies, developmental stages, and cell types is provided in **Supplementary Figures 12-13**. Doublets were filtered out per sample using scDblFinder^110^. Only samples from cortical regions, forebrain, and ganglionic eminences were included, resulting in 2,157,582 cells and 38537 features.

### Cell type annotation

Our cell-type annotation strategy leverages large sets of replicable cell-type markers to perform annotation at the level of individual cells, thereby avoiding the need for cross-dataset integration and cluster-based annotation^31^. This approach uses large gene sets that reproducibly define reference cell types and projects those cell-type annotations onto test datasets using MetaMarkers^31^. Let *x_ij_* be the CPM-normalized expression of gene *i* in cell *j*, and *M_c_* be the set of marker genes for cell type *c*. For each cell *j*, we compute a marker score as:

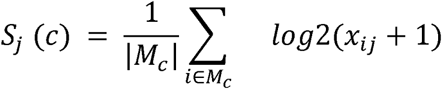

This score is the average marker gene expression of a given marker set for each individual cell, implemented by the MetaMarker score_cells() function. We then compute the marker enrichment score by first computing *s_j_* (*c*) for a series of cell-types *c*_1_,…,*c_n_* and then compute the following for each cell-type:

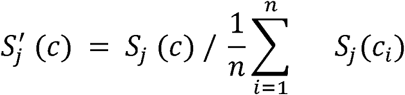

This is implemented by the MetaMarker compute_marker_enrichment() function. This normalizes marker set expression in each cell by the average expression of all marker sets, which corresponds to enrichment under the null that all marker sets are expressed equally. After marker enrichment scores are computed for each cell across all candidate cell-type markers, each cell is assigned the label corresponding to the marker set with the highest enrichment score. This procedure is implemented in the MetaMarkers assign_cells() function.

We used two broad sets of marker genes for annotation: (i) a set of prenatal markers derived across brain regions and datasets spanning first- and second-trimester developmental time points^24^, and (ii) a set of consensus cross-region adult subclass markers^23^. We made several additions and refinements to these initial marker sets.

The prenatal markers were originally computed using the MetaMarkers framework. Briefly, cell-type markers were identified independently within each dataset, and genes were classified as differentially expressed (DE) for a given cell type based on a log2 fold change ≥ 4 and an FDR-adjusted p-value ≤ 0.05. Markers were then ranked across datasets primarily according to the recurrence of differential expression. Thus, highly ranked markers for a cell type are genes that are repeatedly identified as DE across datasets, representing robust and replicable markers of cell-type identity ^24^.

Here, we expanded the original prenatal metamarker collection by adding vascular metamarkers derived from prenatal datasets that included author-supplied annotations for vascular cell types, including Herring^6^ (Vas), Zhu^9^ (Endothelial, Pericytes, and VSMC), and Bhaduri^1^ (Endo and Vascular). We also incorporated marker sets for Cajal-Retzius cells and newborn neurons as defined in ^29^.

The starting set of adult subclass markers was derived from^23^, who provided differential expression statistics for cross-region adult brain single-cell datasets (Jorstad et al. Supplementary Table 4). For each gene, we calculated the average reported log2 fold change across sampled brain regions, requiring representation in at least two regions. Genes were ranked within each subclass according to this average log2 fold change, and the top 100 genes were selected as the initial marker set for each adult subclass.

We further refined these adult marker sets based on the consistency of DE across the Herring^6^, Zhu^9^, and Velmeshev^2^ postnatal datasets. Specifically, postnatal datasets were annotated using the MetaMarkers approach described above with the initial set of 100 adult subclass markers. We then required candidate adult subclass markers to exhibit a cell type-specific fold change ≥ 2 in both the adult dataset^23^ and the earliest sampled post-natal timepoint in at least two of the three postnatal datasets (**Supplementary Fig 14)**. This filtering step identified adult subclass markers with broadly consistent differential expression spanning early post-natal timepoints (0-1 year post-natal) to adulthood.

The final marker sets used for annotation are provided in **Supplementary Table S1**. To generate the cell-type annotations for the prenatal datasets, cells were first annotated using the prenatal metamarker gene sets^24^. For any dataset in the current collection that had been included in the original computation of the prenatal metamarkers, annotation was performed in a leave-one-out manner; see ^24^ for details. A second set of annotations was then generated using the refined postnatal adult marker sets, including the newborn neuron and Cajal-Retzius marker sets.

The final cell-type annotations were obtained by integrating these two annotation schemes. Adult subclass labels were retained unless the corresponding fetal annotation identified the cell as a progenitor population. Finally, cells annotated as newborn neurons were further subdivided into newborn GABAergic or newborn glutamatergic neurons according to their fetal metamarker annotation.

### Joint Embedding

We used a contrastive learning-based approach, CONCORD^111^ to learn the latent space shared by cells and visualize cell type neighborhoods. First, we removed unmapped genes without gene symbols and then identified highly variable genes using the function: ccd.ul.select_features(adata, n_top_features=6000, flavor=’seurat_v3’). Model training used log-normalized data restricted to the highly variable genes, with domain_key set to the concatenation of study, age and region per cell. The UMAP plot was generated using the learned CONCORD embedding and cosine distance with the function: ccd.ul.run_umap(adata, source_key=“Concord”, n_neighbors=30, min_dist=0.1, metric=’cosine’).

### MetaNeighbor

Cell type replicability was assessed using MetaNeighbor’s^25,35^ python implementation (https://github.com/gillislab/pyMN). First, highly variable genes (hvgs) were detected using pymn.variableGenes(adata, study_col=’batch’) with study+age as the batch variable. The anndata object was filtered to these 1318 hvgs for unsupervised all vs all metaneighbor analysis: pymn.MetaNeighborUS(adata, study_col=’batch’, ct_col=’final_annotation’, fast_version=True, symmetric_output=True). Cell type maturation towards adult state was measured by restricting the all vs all matrix to comparisons of all other time points with adult samples for each cell type. The inflection point in cell type maturation trajectories was estimated first with a basic linear fit (lm) to the metaneighbor scores vs age plot, followed by a segmented fit with the breakpoint initialized as the median age.

Cell type replicability between developing human and mouse brains were estimated using the developing mouse visual cortex dataset from the Allen Institute^17^. We converted mouse cell type nomenclature to match our human atlas (e.g. “Pvalb Gaba” ∼ “Pvalb”, “L6 CT CTX Glut” ∼ “L6 CT” etc.). Human and mouse datasets were combined and restricted to 15638 one-to-one orthologs and highly variable genes were identified using species as the batch variable. 1631 hvgs were used to run unsupervised all vs all MetaNeighbor analysis.

### Spatial Projection

We use tangram^40^ to project single-cell annotations from our jointly processed atlas into MERFISH spatial transcriptomics data from the developing human cortex^26^. Each cell in the spatial data is assigned the cell type with highest probability Tangram prediction. Keeping the ventricular zone (vz) as the reference, we computed relative distance of each cell from the nearest vz cell (cKDTree from scipy.spatial) to determine relative cortical depth of different cell types. Enrichment of cell types in author-defined cortical zones was computed using the hypergeometric test and P-values across individual samples were aggregated using Fisher’s method.

### Cell type-specific maturation models

Gene expression data from prenatal samples were aggregated by cell type and sample using Seurat’s AggregateExpression, resulting in a total of 3542 samples, and features were restricted to protein-coding genes expressed in atleast 10% of the samples (N = 16,373). A meta-analytic regularized regression model (“glmnet”, caret package) was trained to use log-normalized gene expression to predict age within each cell type. Importance scores for genes used by the models were computed using two approaches: 1) bootstrap importance scores were estimated by retraining 100 models per cell type on random resamples of the training data and calculating gene usage frequency, 2) Perturbation importance scores were estimated by randomly permuting the expression of each gene across the samples and evaluating the percentage drop in model performance (mean absolute error). Both importance scores and model coefficients for all cell types are reported in **Supplementary table S3.**

To identify top cell type-specific maturation genes, we first computed spearman correlation between age and gene expression for all genes within each cell type, and then computed a specificity score^112^ (*τ*) to quantify cell type specificity of age-correlations:

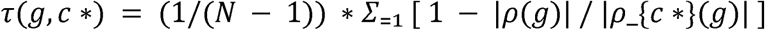

where ρ (g) is the age-correlation of gene *g* in cell type *i*, c* is the focal cell type, and N is the number of cell types evaluated. Values near 1 indicate that the correlation magnitude is concentrated in c*, values near 0 indicate comparable correlation magnitudes across cell types, and negative values indicate that one or more non-focal cell types have stronger age-correlations than c*. Genes were ranked by high bootstrap importance scores and filtered to those with *τ*(*g*,*c*\*) > 0.5. Top 50 genes per cell type and their importance and tau scores are provided in **Supplementary table S4.**

### Microglia trajectory analysis

35111 Immune cells were subclustered using CONCORD embeddings optimized by the MrTree package in R and cell type and states were annotated based on established markers and donor ages. The trajectory of microglial states was constructed using the Monocle3 package and rooted in the Mic - nascent cluster, which had the highest expression of progenitor markers. Differentially expressed genes for the branches were calculated using a Moran’s I test within the Monocle3 framework.

The Yolk Sac single cell RNA seq data was downloaded from https://developmental.cellatlas.io/yolk-sac and the MetaNeighbor algorithm was used to determine AUROC in R with 3299 hvgs across the two datasets.

Age trends were determined by calculating the log (base 10) of the proportion of microglial cell states per donor and a generalised additive model was used to fit the log(proportion) with the age (years). The curves were plotted with LOESS smoothing in ggplot2 in R.

### NMF Pattern Projections

NMF pattern loadings were obtained from a supplementary table in Sonthalia et. al. The projectR package was used to project each set of patterns into 200,000 cell subsets of the complete log normalised dataset.

### Meta-module generation

Meta-modules characterizing human brain development before GW 14 and between GW 20 – 25 were generated as previously described^29^ and provided in **Supplementary tables S11-12**. For computational efficiency, the counts matrices for several individual samples were subsetted prior to meta-module generation. These include the subsetting of the under GW 14 samples CS22_10 (Eze, et al dataset) and NEOCORTEX_13 (Salamon, et al dataset) into 3 subsets; the GW 20 – 25 samples GW 34_20 (Bhaduri et al., dataset) and NEOCORTEX_20 (Salamon, et al dataset) into 3 subsets; and the GW 20 – 25 samples GW22_22, GW20_31_20, and GW20_20 (Bhaduri et al., dataset) into 4 subsets.

### Co-expression analysis of metamodules

We constructed two complementary co-expression networks to evaluate module co-expression. For the **across-cell-type network,** we pseudobulked cells by cell type within each study, computed gene–gene Spearman correlations, then rank-standardized matrix entries to [0,1] and aggregated study-specific matrices by averaging the rank-standardized entries. Variation in this aggregate network is dominated by cross-cell-type mean differences.

For the **within-cell-type-network,** we sampled 100 random pseudobulks of 10 cells from each cell type × study, computed and rank-standardized gene–gene correlations within study, and averaged across studies to obtain an aggregate co-expression network for each cell type. We then averaged the per-cell-type networks across cell types. Because each input network is constructed from cells of a single type, cross-cell-type mean differences are excluded by construction and the final aggregate reflects average within-cell-type co-expression.

Overall, we generated across- and within-cell type co-expression networks for four age windows: first trimester (< GW14), second trimester (GW 14-25), third trimester - neonatal (> GW33 - < 1 year), and postnatal (> 1 year old) stages.

Module co-expression strength in each network was scored with EGAD^83^ using three-fold neighbor-voting cross-validation: module genes are partitioned into folds, and for each held-out fold, membership is predicted from the co-expression network of the remaining module genes. The resulting AUROC indicates how tightly the module’s genes are co-expressed in that network.

### Software and code

Key results are available to explore in <u>brain-development browser</u> and our data will be made available for download at CellxGene and Zenodo (https://doi.org/10.5281/zenodo.21375950). All code and data required to reproduce key figures are available on Github (https://github.com/sridevi96/DevBrainAtlas) and Zenodo.

## Acknowledgements

This work was supported by the National Institutes of Health: U24 MH130968 to JG; R24 MH114788 (NeMO Archive), U24 MH130968 (CUBIE), R01 MH052716 to SAA, CC, BH, and SM; R01 MH132689, UM1 MH130991 to AB; U01 MH130962, UM1 MH130991, R01 MH125516 to TN; Natural Sciences and Engineering Research Council of Canada: NSERC Discovery RGPIN-2025-06198 to JG; Mclaughlin Centre Accelerator Grant to SV and JG; SV was supported by the Canadian Institutes of Health Research Fellowship and Schmidt Science Fellows, in partnership with the Rhodes Trust; LW was supported by NIMH R00MH131832, the Stinehart-Reed Award, and the Virginia and D.K. Ludwig Fund for Cancer Research.

We thank members of the BICAN Developmental working group and the BICAN developmental joint analysis subgroup for their helpful feedback.

