## Supplementary Figures for "A consensus atlas of human brain development defines cell type-specific maturation trajectories across the lifespan"

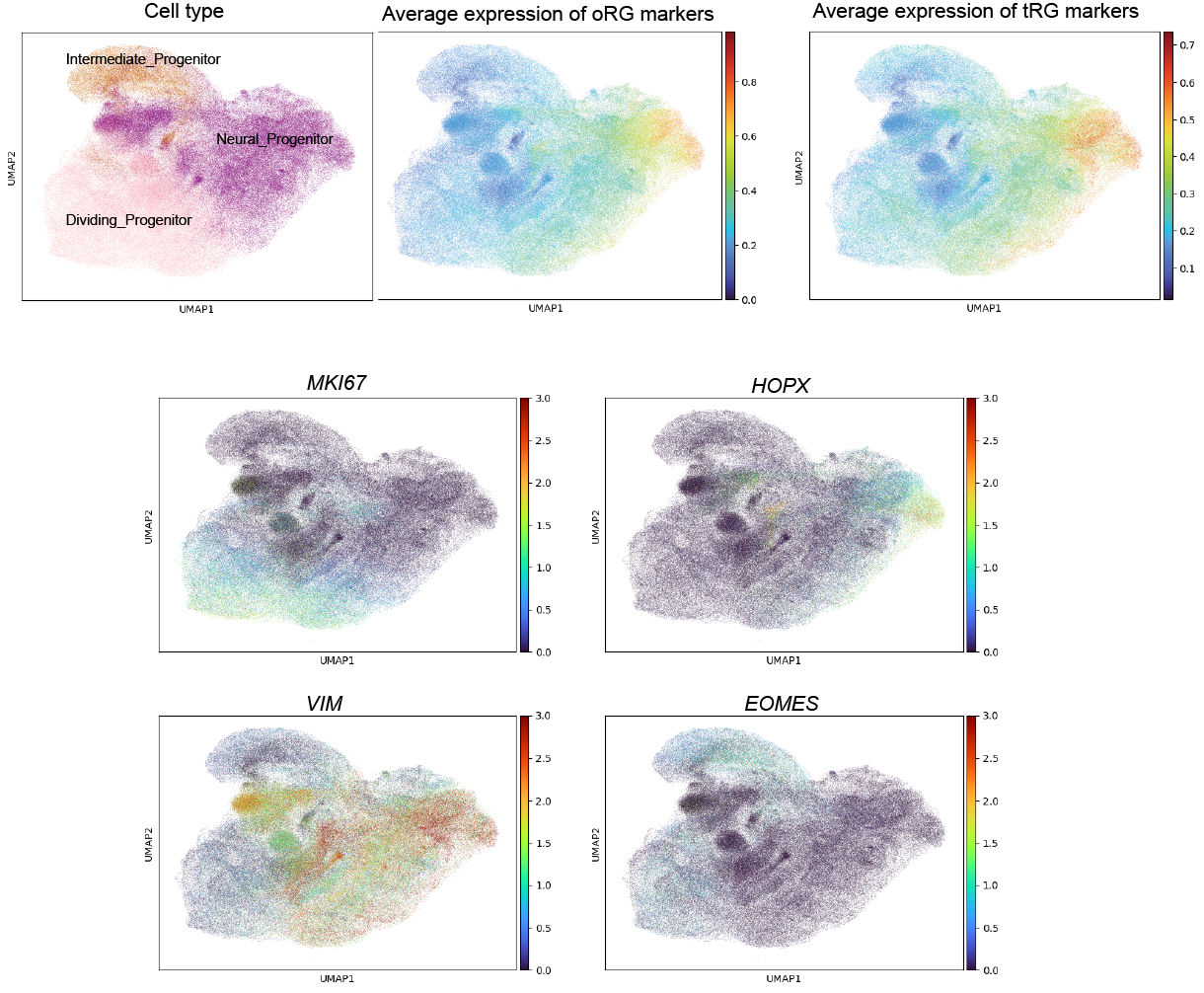

**Supplementary Fig 1: Radial glia marker expression in Progenitor subclasses**

(Top) UMAP of progenitor cell types, with average expression of outer radial glia (oRG) and truncated radial glia (tRG) marker gene sets from Nano et al. 2025. (Bottom) UMAP showing expression of canonical progenitor marker genes: *MKI67* (Dividing Progenitors), *HOPX* (oRG), *VIM* (pan-RG), *EOMES* (Intermediate Progenitors).

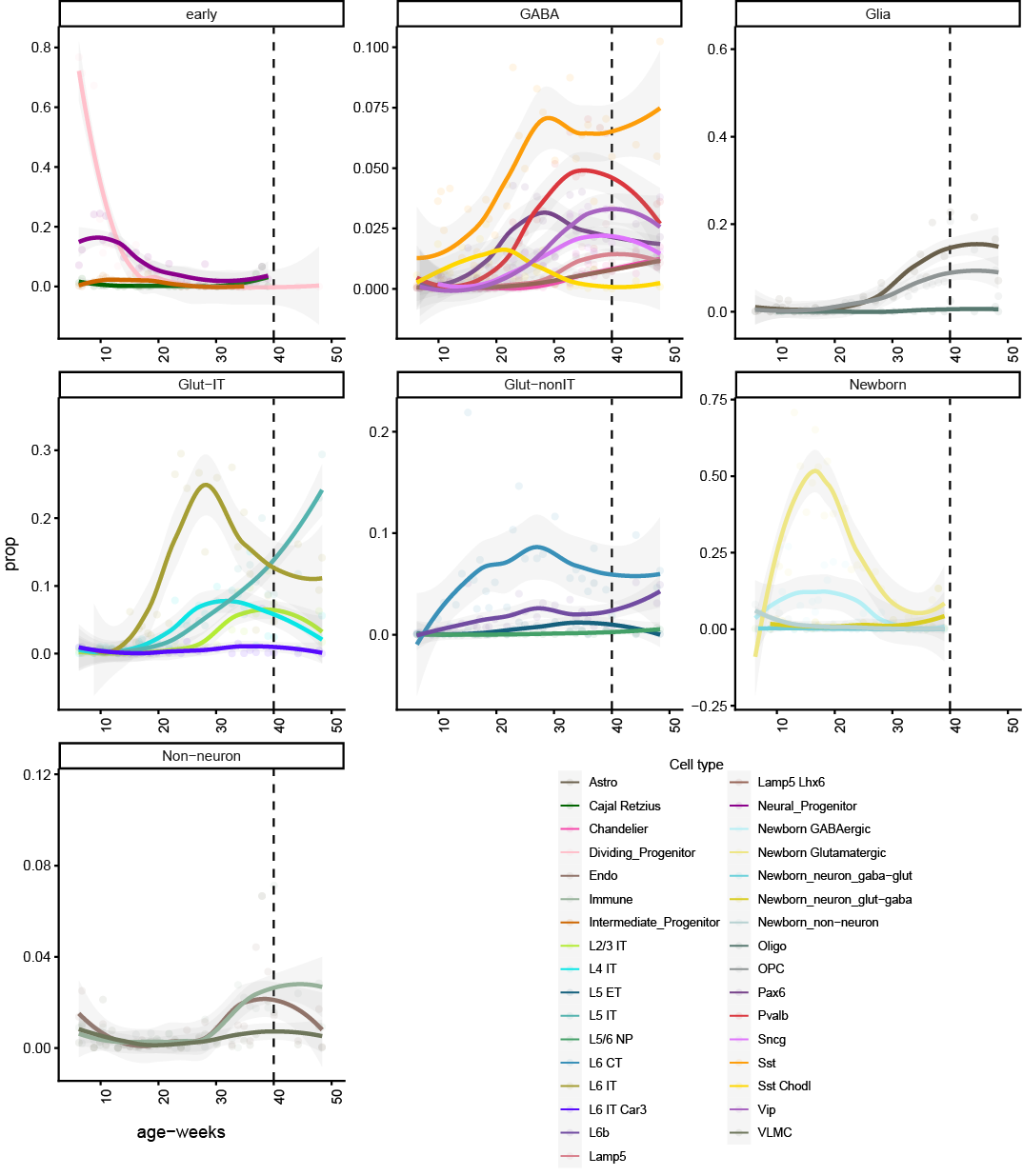

**Supplementary Fig 2: Changing cell type proportions over age**

Panels show smoothed fits to proportions of various cell types present at different timepoints in the prenatal to perinatal periods in our jointly processed atlas

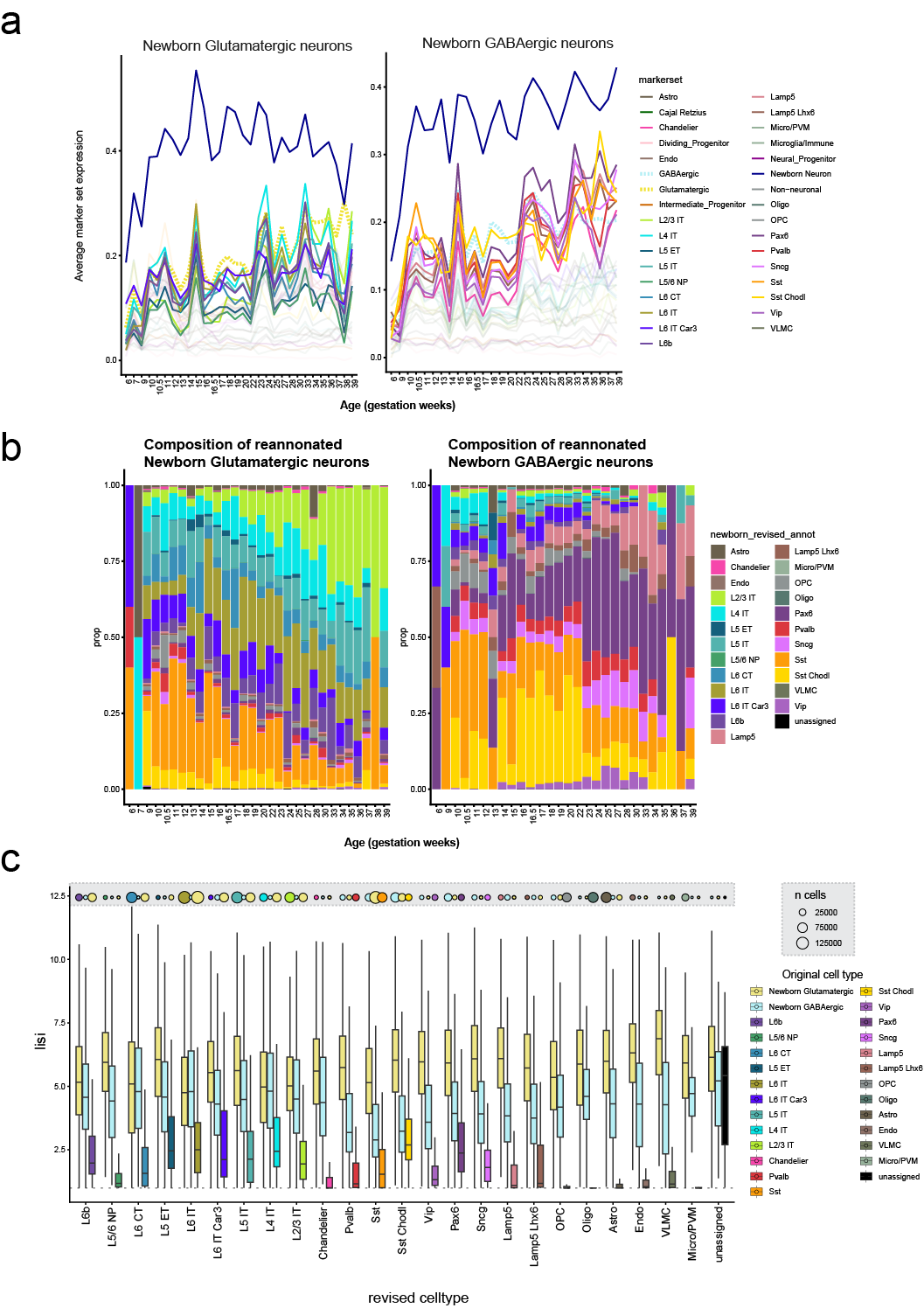

**Supplementary Fig 3: Biased marker expression and diversity of newborn neurons**

**a,** Expression of newborn neuron and specific subclass marker sets in newborn glutamatergic (left) and GABAergic (right) neurons. **b,** Newborn glutamatergic/GABAergic neurons were reannotated as specific subclasses based on their next highest marker enrichment. Graphs show proportions of re-annotated newborn neuron subclasses across age. Newborn neurons are biased towards different subclasses at different timepoints. **c,** LISI scores computed from 2D UMAP for newborn neurons using their original newborn annotation or re-annotated subclass annotation, compared to the LISI scores of corresponding mature subclasses. Newborn neurons have generally high LISI scores indicating the heterogeneity of their cell neighborhoods in the UMAP.

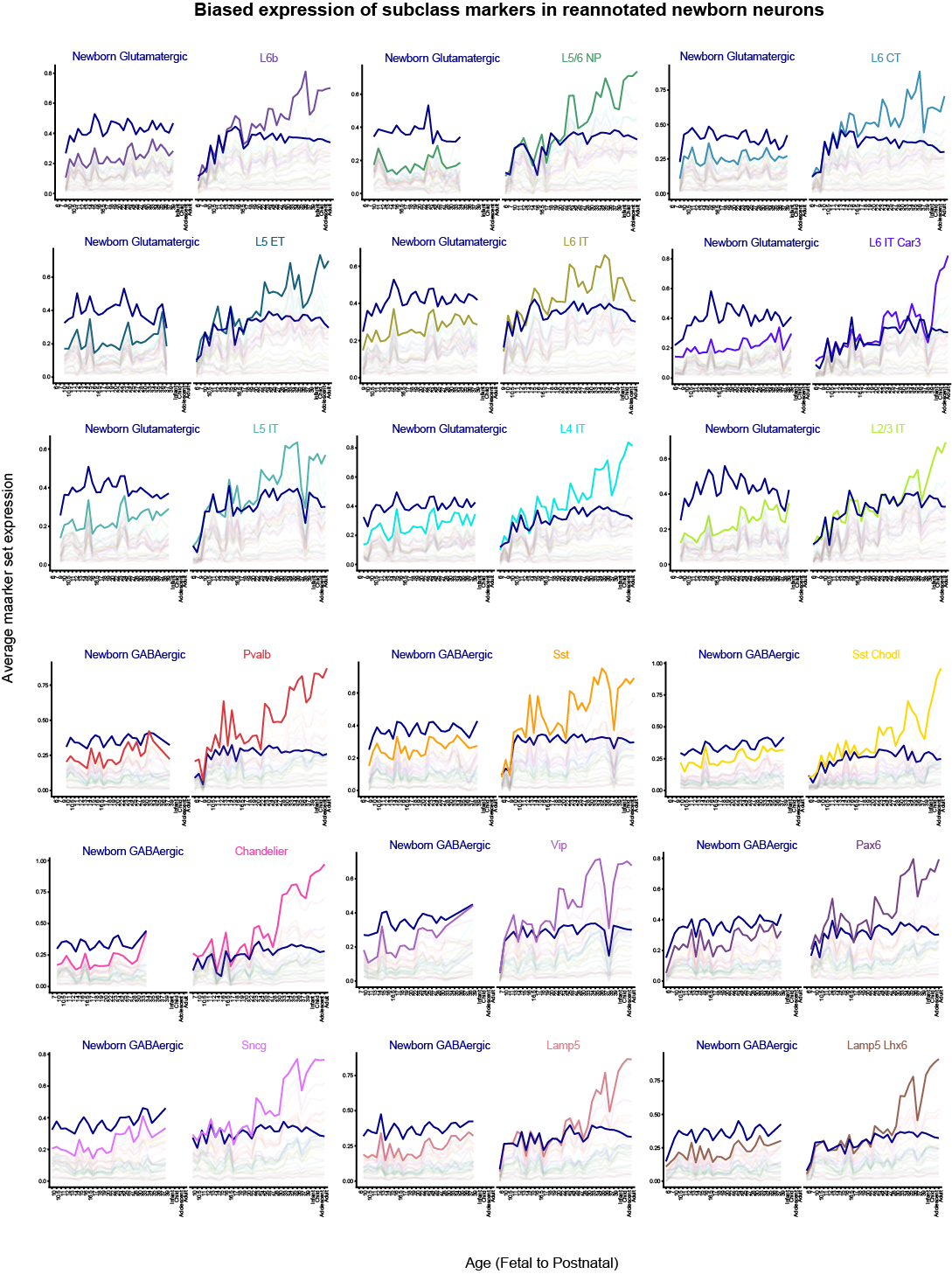

**Supplementary Fig 4: Marker expression in re-annotated newborn neurons**

Each panel shows average expression of specific subclass markers in re-annotated newborn neurons (left) and the same marker set’s expression in the corresponding mature subclass (right) over age (x-axis).

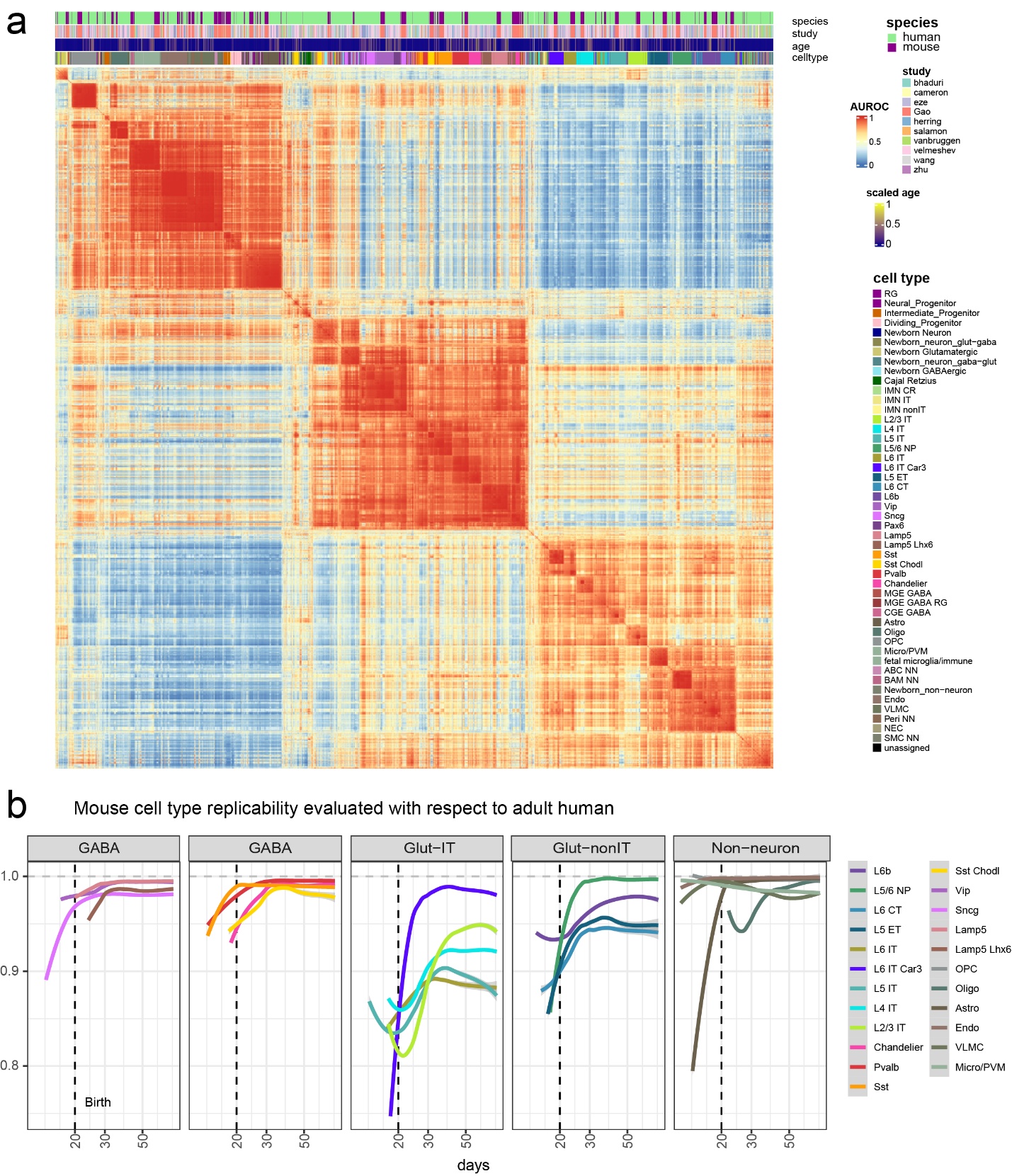

**Supplementary Fig 5: Comparing developing cell types in the human and mouse brain**

**a,** Heatmap of MetaNeighbor scores measuring cell type replicability between the mouse and human brain across time points and samples. **b,** MetaNeighbor scores for mouse cell types at different timepoints with respect to the adult human samples are plotted.

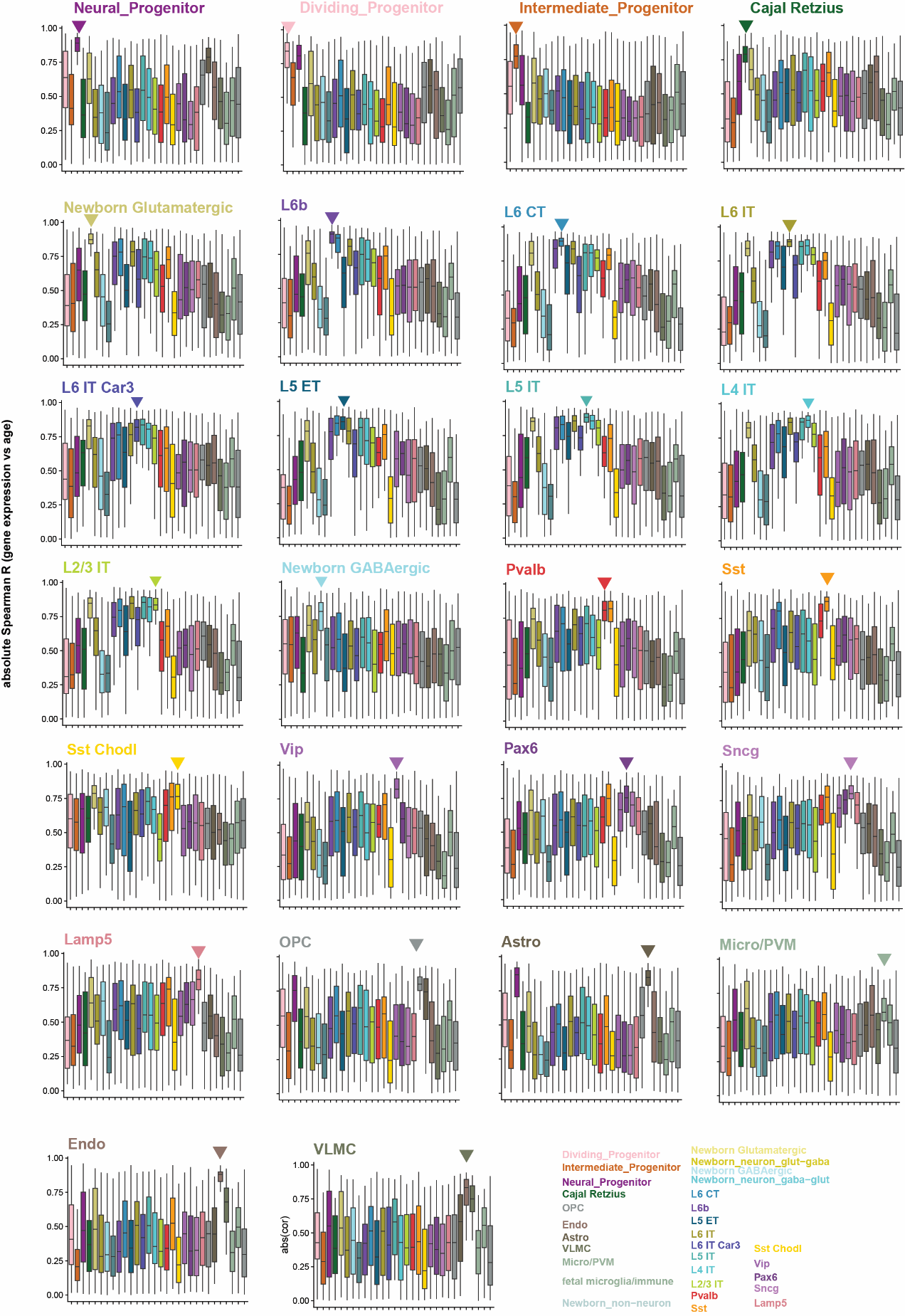

**Supplementary Fig 6: Age-correlation of top 50 cell type-specific maturation genes**

Each panel shows the distribution of absolute spearman correlation with age for the top 50 maturation genes for that cell type, with x axis ordered by the cell type with highest median correlation. Cell type-specific genes have highest correlation to age in that cell type.

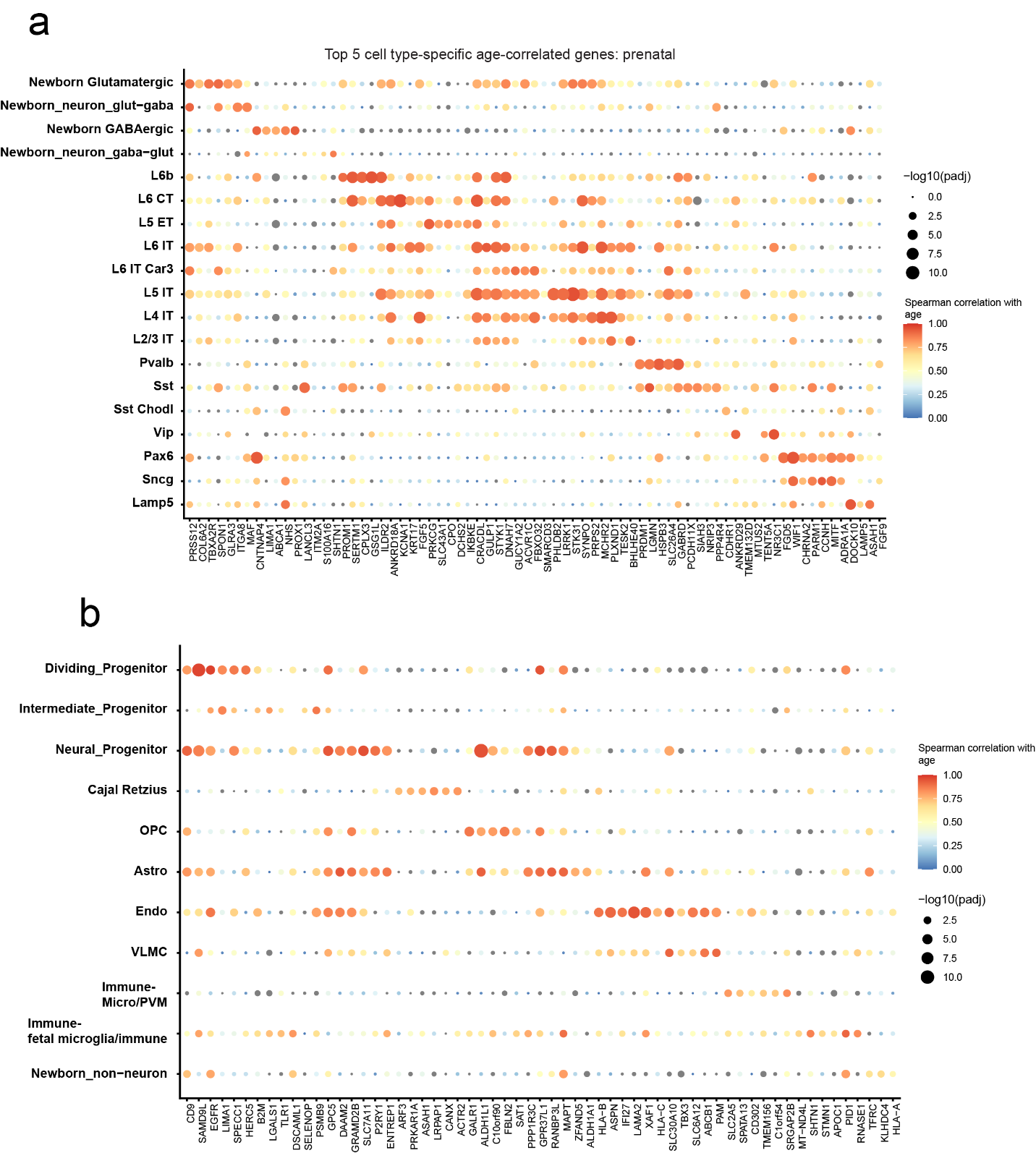

**Supplementary Fig 7: Top 5 positive cell type-specific maturation genes**

Dot plots show correlation of top 5 specific genes per cell type with age. Dot size indicates -log10(FDR adjusted P-value) and color indicates the correlation coefficient in **a,** Neurons and **b**, progenitors and non-neurons

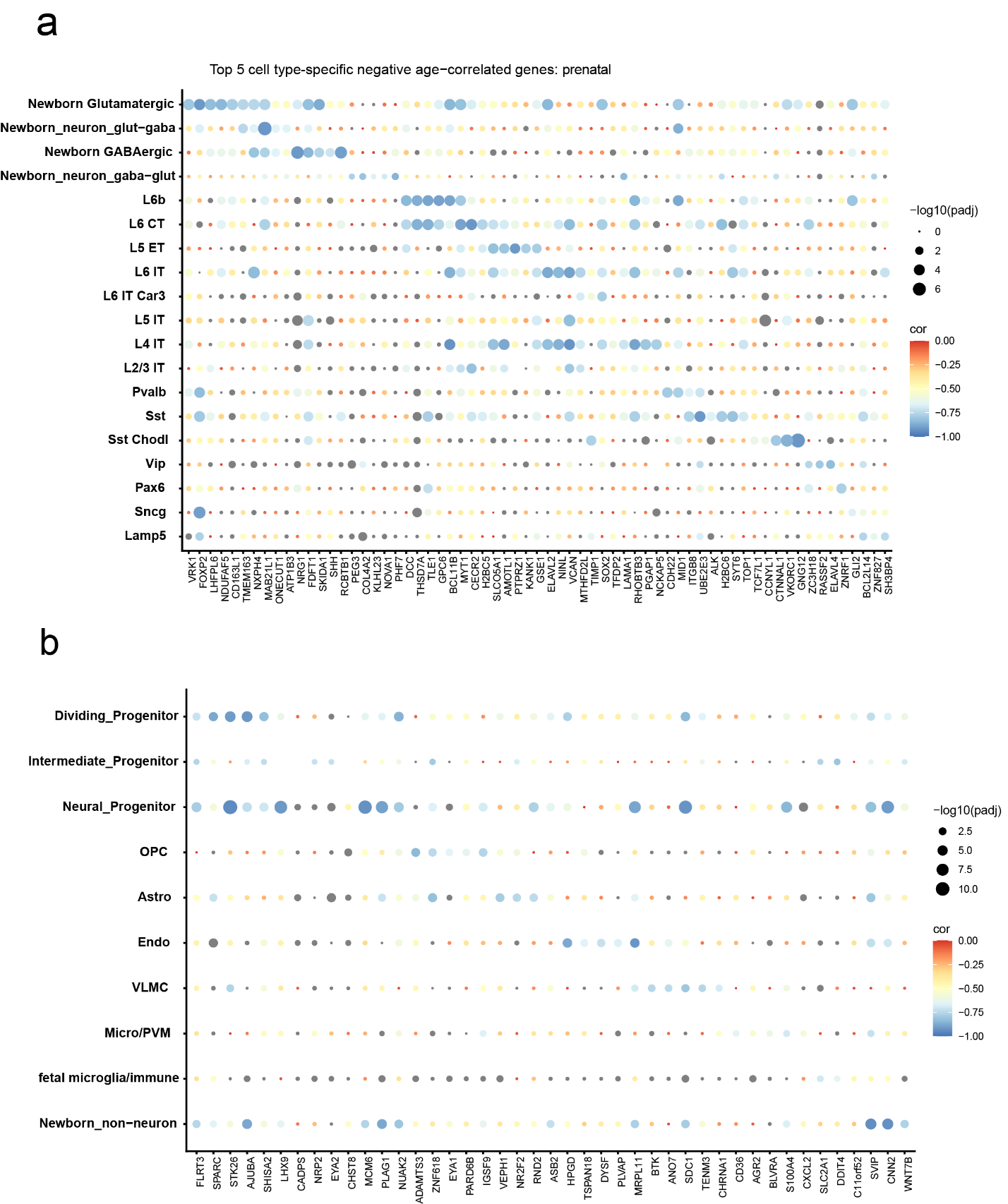

**Supplementary Fig 8: Top 5 negative cell type-specific maturation genes**

Dot plots show correlation of top 5 specific genes per cell type with age. Dot size indicates -log10(FDR adjusted P-value) and color indicates the correlation coefficient in **a,** Neurons and **b**, progenitors and non-neurons

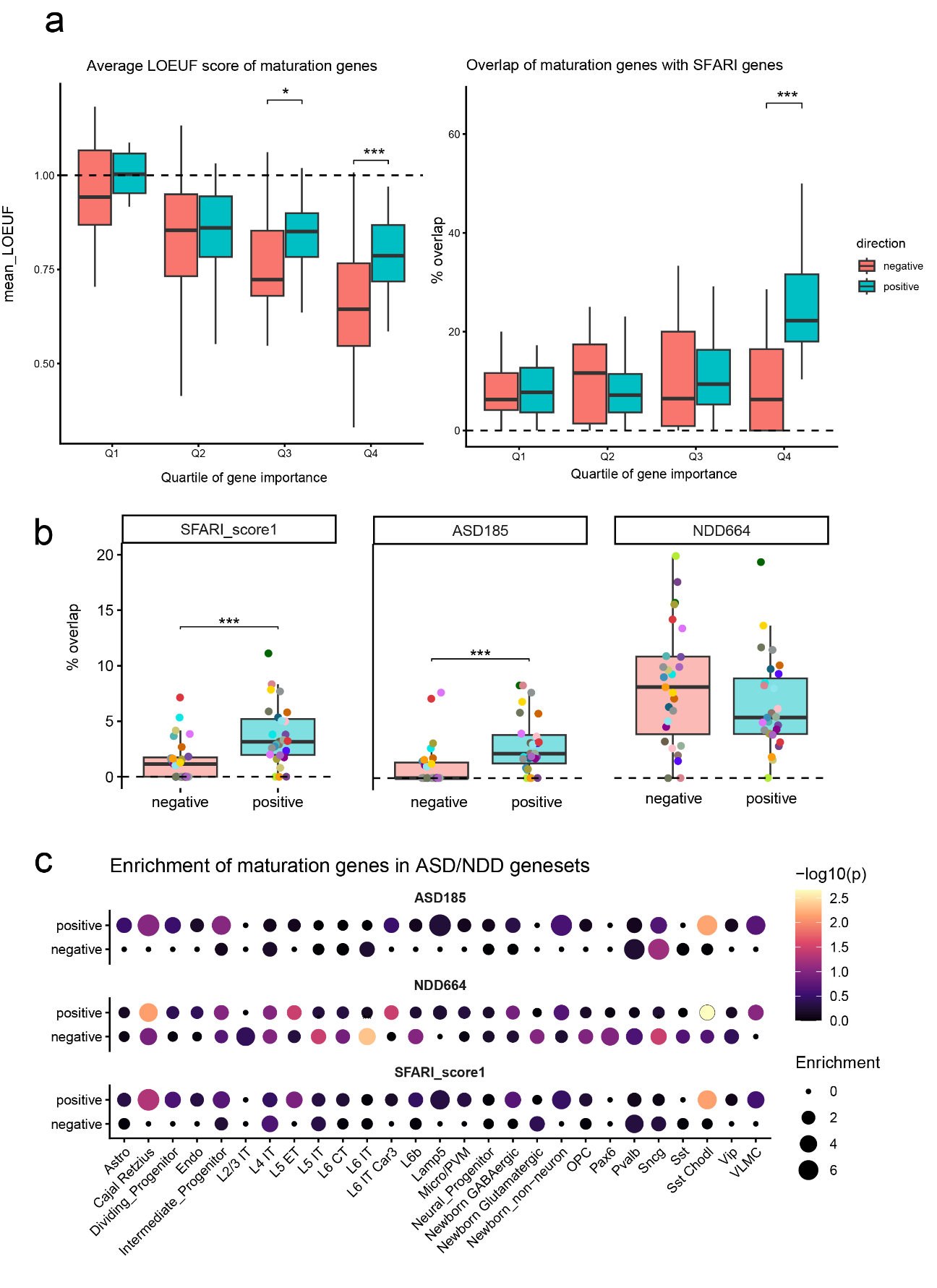
**Supplementary Fig 9: Evolutionary conservation and disease overlap of cell type-specific maturation genes**

**a**, **Left**- Binned distribution of LOEUF scores for positive (upregulated) and negative (downregulated) genes at different quartiles of their bootstrap importance scores (x-axis). **Right-** Binned distribution of percentage overlap with SFARI genes for positive (upregulated) and negative (downregulated) genes at different quartiles of their bootstrap importance scores (x-axis). * P< 0.05, ** P<0.01, ***P<0.001 FDR adjusted Wilcoxon test. **b**, Percentage overlap of each cell type’s positive and negative maturation genes with neurodevelopmental disorder (NDD) rare variant genes: SFARI score 1, ASD185 and NDD664 genes from Fu et al. 2022. c, Dot plot shows enrichment of disease gene sets in cell type-specific maturation genes.

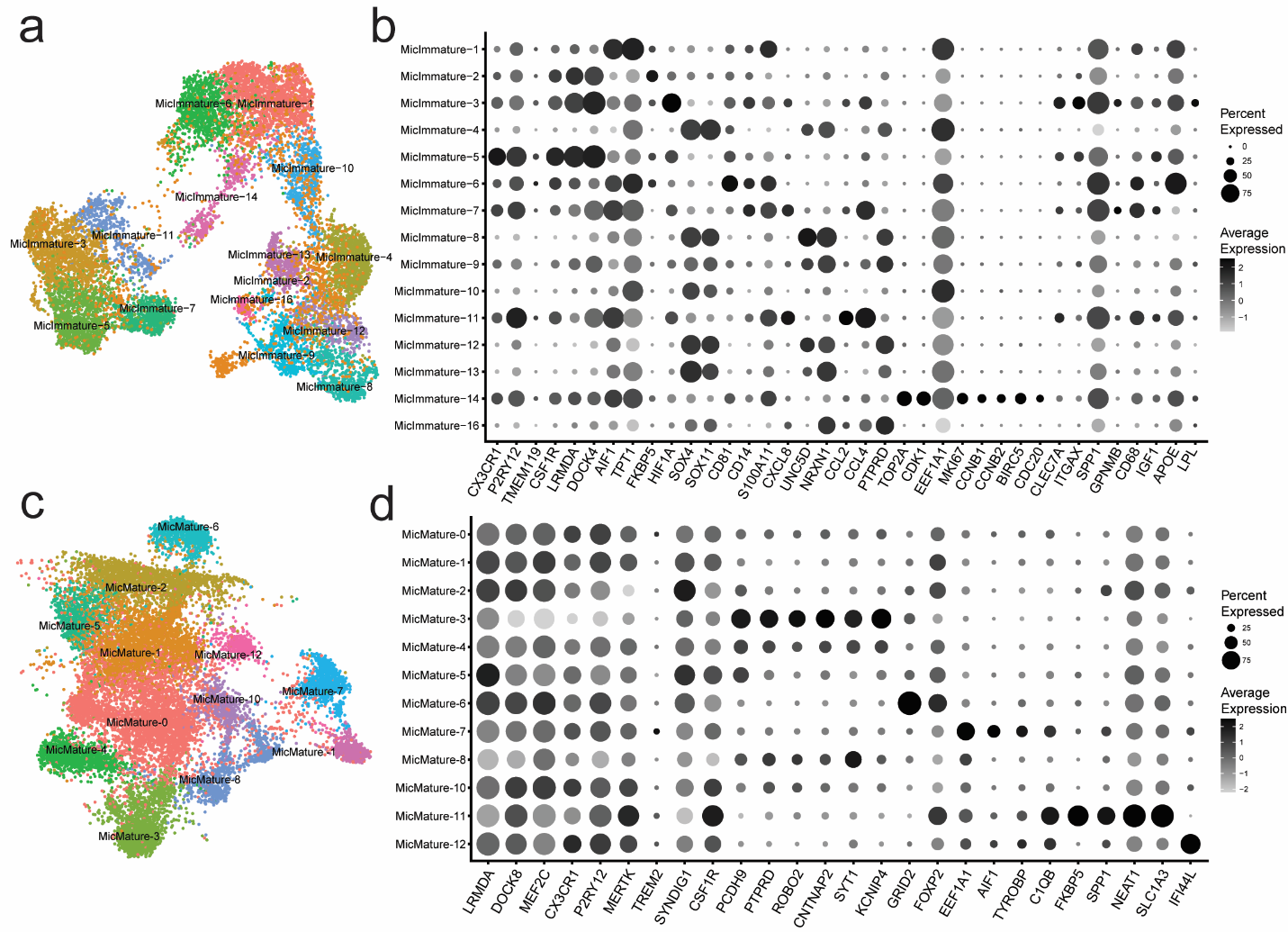

**Supplementary Fig 10: Exploring the diversity of immature and mature microglia**.

a) UMAP of the subclusters of immature microglia. b) Dotplot displaying the expression of marker genes for each subcluster of the immature microglia. c-d, Same plots for subclusters of mature microglia

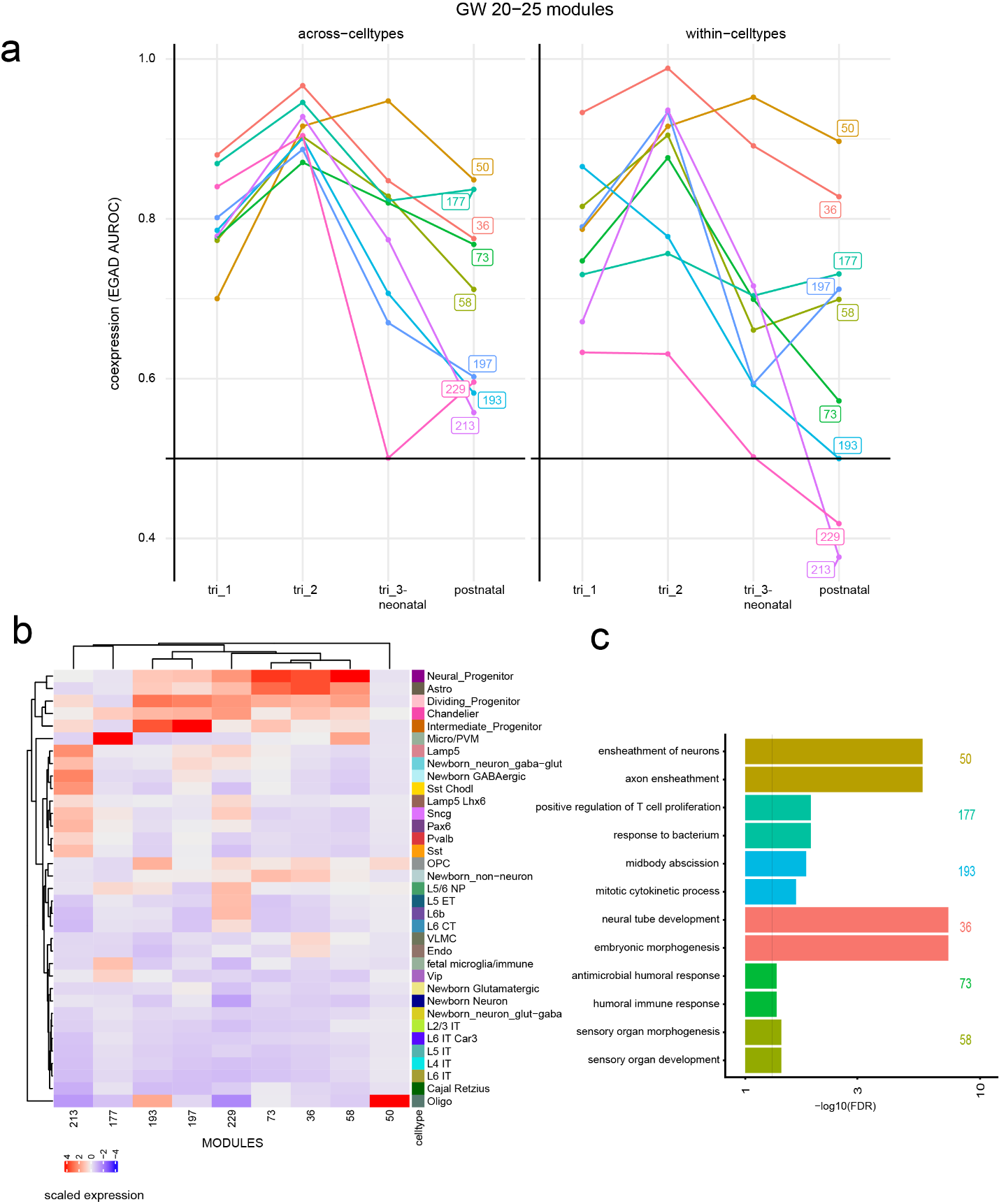

**Supplementary Fig 11: Properties of selected gene modules from mid-gestation**

**a,** Co-expression (EGAD AUROC) for select modules from GW20-25 at different time points across (left) and within (right) cell types. **b**, Module expression in specific cell types. **c**, GO enrichment for selected modules.

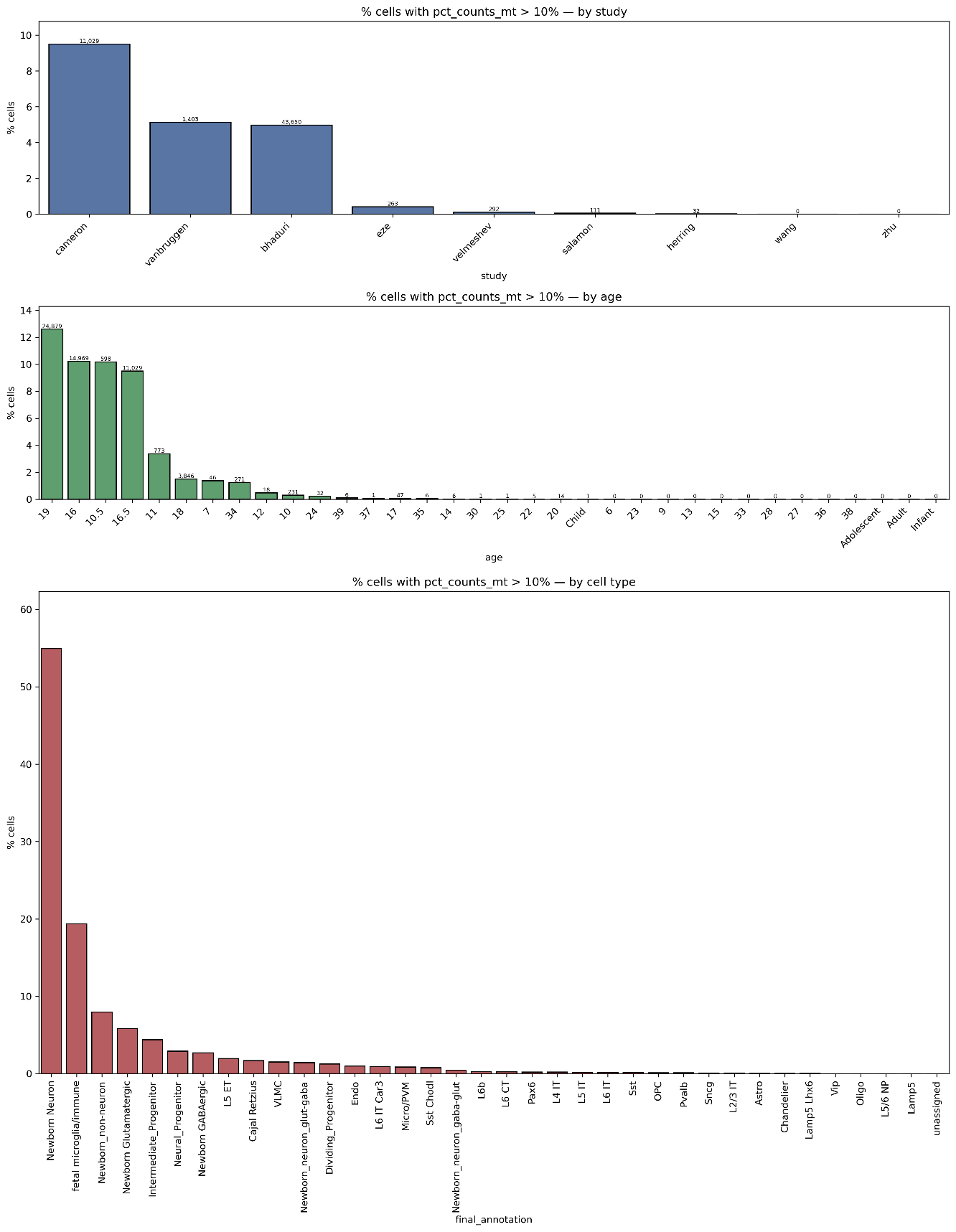

**Supplementary Fig 12: Mitochondrial counts distribution**

Percentage of cells per study (top), age (middle) and cell type (bottom) with > 10 % mitochondrial counts. Most datasets have fewer than 5% of cells with high mitochondrial counts. ‘Newborn Neuron’ which has the highest proportion of cells with high mitochondrial counts is a small set of ambiguous 131 cells which were not successfully mapped to either Glutamatergic or GABAergic identity.

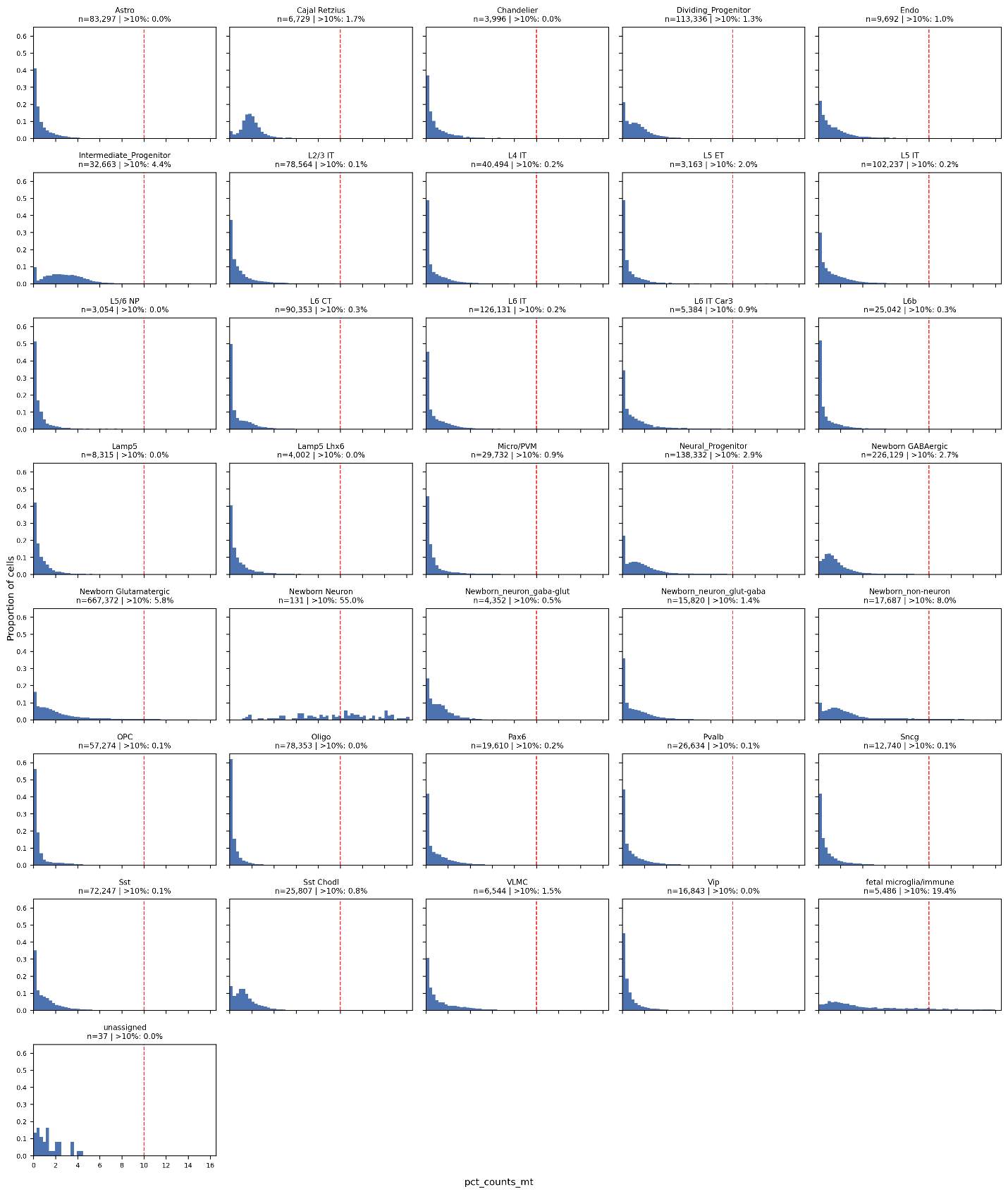

**Supplementary Fig 13: Mitochondrial percentage distribution per cell type**

Histograms show distribution of percentage of mitochondrial counts in each cell type. Red dotted line denotes 10% threshold. Only ‘Newborn Neuron’ cells show high % mitochondrial counts beyond the threshold. These cells were not used for downstream analyses.

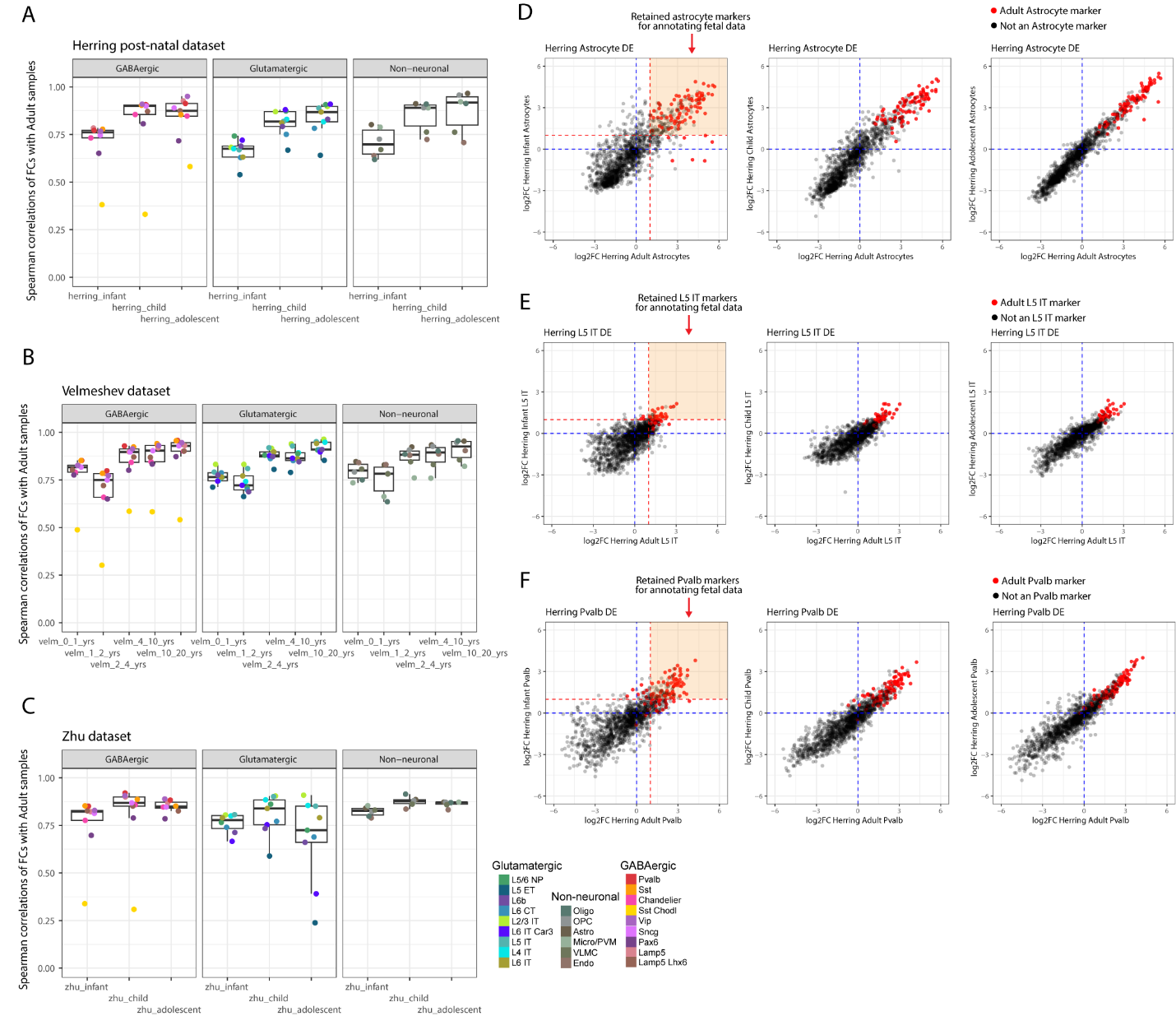

**Supplementary Fig 14: Marker gene set refinement based on consistent differential expression across time points**

**a**. Boxplots depicting the spearman correlations of subclass fold-changes (FCs) for post-natal time points compared to adult data in the Herring post-natal dataset. Panels D, E, and F provide several examples of the gene-level FC comparisons used to compute these correlations.
**b**. Same as A, but for the Velmeshev post-natal dataset.
**c**. Same as A, but for the Zhu post-natal dataset.
**d**. Scatter plot of gene expression FCs of all genes computed for Astrocytes vs all other cell-types in the Herring post-natal dataset. Left-most plot compares infant FCs to adult FCs, middle plot compares child FCs to adult FCs, and right-most plot compares adolescent FCs to adult FCs. The starting 100 adult astrocyte markers are colored red. The orange box highlights the adult markers that have FCs >= 2 in both the adult data and the infant data. FCs are log2 transformed in the plots.
**e**. Same as D, but for L5 IT expression and markers.
**f**. Same as D, but for Pvalb expression and markers.
